# Endogenous formaldehyde scavenges cellular glutathione resulting in cytotoxic redox disruption

**DOI:** 10.1101/2020.05.14.090738

**Authors:** Carla Umansky, Agustín Morellato, Marco Scheidegger, Matthias Rieckher, Manuela R. Martinefski, Gabriela A. Fernandez, Ksenia Kolesnikova, Anna J. Vesting, Ismene Karakasilioti, Hernán Reingruber, Yan Wei, Rongqiao He, Mariela Bollini, María Eugenia Monge, Björn Schumacher, Lucas B. Pontel

## Abstract

Formaldehyde (FA) is a ubiquitous endogenous and environmental metabolite that is thought to exert cytotoxicity through DNA and DNA-protein crosslinking. We show here that FA can cause cellular damage beyond genotoxicity by triggering oxidative stress, which is prevented by the enzyme alcohol dehydrogenase 5 (ADH5/GSNOR). Mechanistically, we determine that endogenous FA reacts with the redox-active thiol group of glutathione (GSH) forming S-hydroxymethyl-GSH, which is metabolized by ADH5 yielding reduced GSH thus preventing redox disruption. We identify the *ADH5*-ortholog gene in *Caenorhabditis elegans* and show that oxidative stress also underlies FA toxicity in nematodes. Moreover, we show that endogenous GSH can protect cells lacking the Fanconi Anemia DNA repair pathway from FA, which might have broad implications for Fanconi Anemia patients and for healthy *BRCA2*-mutation carriers. We thus establish a highly conserved mechanism through which endogenous FA disrupts the GSH-regulated cellular redox homeostasis that is critical during development and aging.

## Introduction

FA is a potent genotoxin classified by the World Health Organization (WHO) as a human carcinogen^1^. In the body, FA can originate from cellular metabolism, i.e. histone and DNA demethylation reactions or the one-carbon cycle; it can also arise from the diet and it is ubiquitously found in the environment^2–5^. Indeed, this aldehyde is more abundant in the body than previously thought;different works have reported FA quantification in healthy human blood samples with levels in the 10-50 μM range^6,7^. Endogenous FA has been suggested as a causative agent for several human diseases such as Fanconi Anemia and the Ruij-Aalfs syndrome, and it might drive cancer in *BRCA2*-mutation carriers^5,8^. Indeed, Fanconi Anemia patients carrying a mutation in the acetaldehyde/FA metabolizing gene *ALDH2* present accelerated progression of bone marrow failure (BMF)^9–11^. Moreover, mice lacking *ADH5* and the Fanconi Anemia DNA repair pathway show severe BMF, liver and kidney dysfunction and early cancer onset^9^, indicating that endogenous FA can drive cancer initiation and Fanconi Anemia phenotypes.

Genotoxicity has been widely indicated as the main consequence of FA reactivity in cells^4^. However, the strong reactivity of the FA carbonyl group might also affect other molecules than DNA. *In vitro*, the spontaneous electrophilic attack of the FA carbonyl group to the thiol-group of GSH leads to the formation of the covalent product S-hydroxymethyl-GSH (HSMGSH)^12^. This reaction might be strongly favored inside cells, where GSH levels are in the millimolar range^13^. Accordingly, ADH5 metabolizes HSMGSH yielding formate, which is directed to the one-carbon cycle for nucleotide synthesis^3^ (Fig. 1A).

**Fig. 1.**
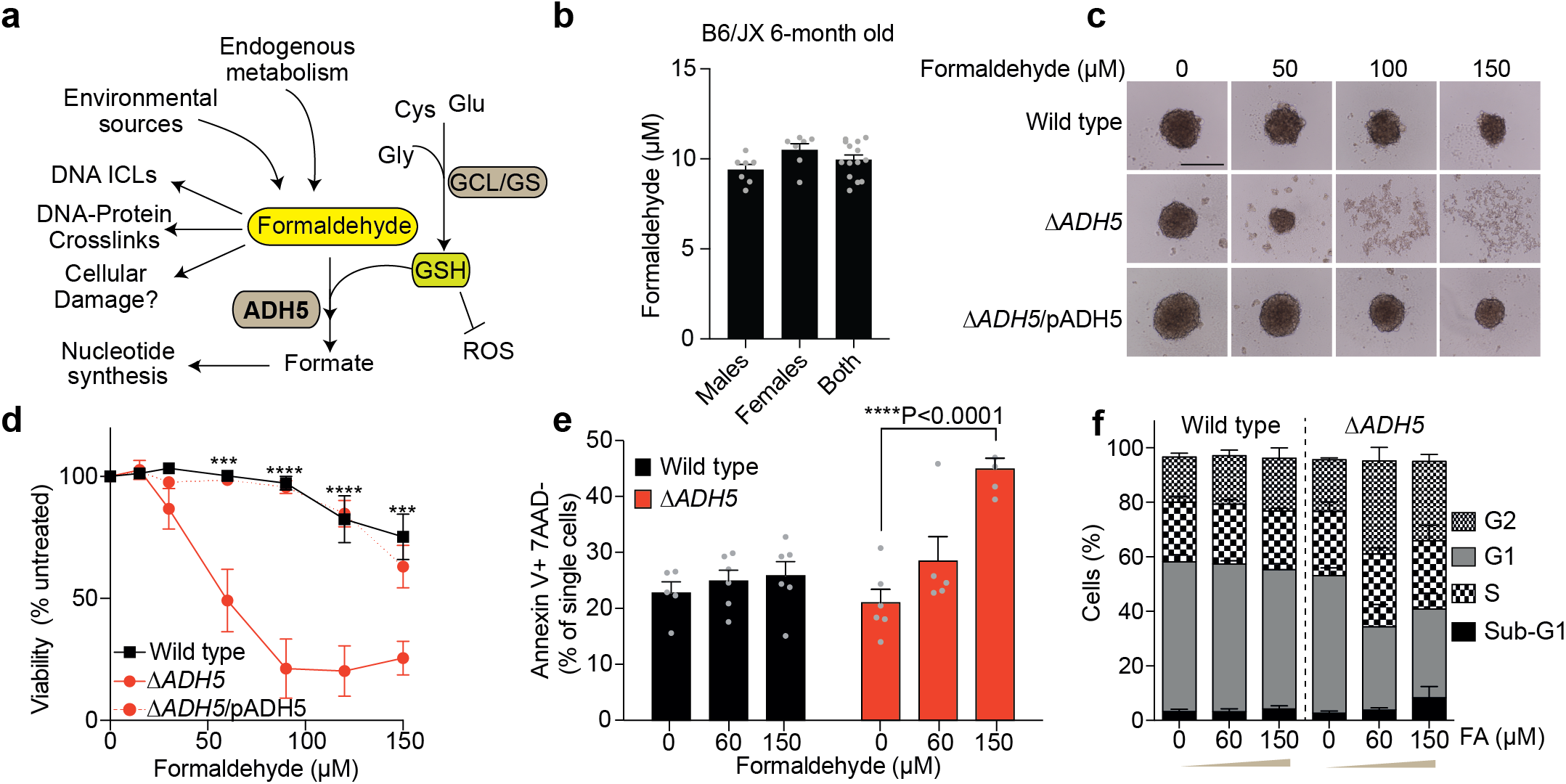
ADH5 prevents formaldehyde toxicity in human cancer cells. **a.** Scheme showing formaldehyde (FA) metabolism, glutathione (GSH) *de novo* synthesis and the central role of ADH5 (alcohol dehydrogenase 5). GCL: glutamate cysteine ligase; GS: glutathione synthetase; ROS: Reactive Oxygen Species. **b.** FA in blood of 6-month old mice determined by high-performance liquid chromatography with ultraviolet detection (UV-HPLC) (mean ± s.e.m., n = 7 (males), n = 7 (females), n = 14 (both sexes). **c.** Representative images of 3D-sphere formation for HCT116 Wild type (WT), Δ*ADH5* and Δ*ADH5*/*p*ADH5 cells (scale bar 0.5 mm). **d.** Resazurin-based viability assay expressed as % of fluorescence relative to untreated cells. Each point represents the mean ± s.e.m. of 6 independent experiments done by triplicate. Asterisks represent the statistical significance according to one-way ANOVA for multiple comparison using a Tukey-corrected test between Wild type and Δ*ADH5* cells. **e.** Apoptosis determined by Annexin V detection in cells exposed to FA over 24 h at the concentrations indicated in the figure (mean ± s.e.m., n=5, unpaired t-test). **f.** Cell cycle analysis of Wild type and Δ*ADH5* HCT116 cells exposed to 60 and 150 μM FA over 24 h (mean ± s.e.m., n = 4).

Considering the electrophilicity of FA and the abundance of GSH we hypothesize that the reaction between FA and GSH might affect the GSH pool having detrimental biological consequences. Indeed, alterations in GSH homeostasis have been reported in multiple pathologies such as hemolytic anemia, diabetes, liver diseases, cystic fibrosis, neurodegeneration and cancer^14–17^. GSH not only neutralizes reactive oxygen species (ROS), but can also promote chemoresistance by forming GSH-xenobiotic conjugates that are pumped out of the cell *via* multiple resistance-associated protein transporters (MRP)^18^. To replenish intracellular GSH, cells synthesize GSH in a two-step metabolic pathway centered on the rate-limiting enzyme glutamate cysteine ligase (GCL), which is composed of a catalytic (GCLC) and a regulatory (GCLM) subunit, and the GSH synthetase (GS) (**Fig. 1a**)^18^. Cells might thus also need to maintain the balance between GSH and the oxidized GSH disulfide form (GSSG) -GSH:GSSG-to limit free FA and to prevent redox disruption.

We report here that FA toxicity is inflicted by the reaction between FA and the redox-active thiol group present in GSH, which disables the antioxidant property of GSH. Our data also support a previously unrecognized function of GSH in the protection against FA toxicity, and an evolutionary conserved mechanism that maintains GSH:GSSG balance by salvaging reduced GSH from FA-GSH covalent adducts. These data might have wide implications not only for Fanconi Anemia patients and *BRCA2*-mutation carriers but also for cancer cells that would have to overcome blood FA level for a successful disease progression^8,9^.

## Results

### ADH5 prevents FA toxicity in human cancer cells

FA levels in blood from different species have been reported in the 10-50 μM range (Reingruber and Pontel, 2018 and references therein). With the aim to determine the amount of FA in blood, we set out to measure this aldehyde in serum samples from 6-month old mice. FA was detected and quantified in mouse blood with a mean concentration of 9.95 ± 1 μM (n=14), which is in the same order as values reported for healthy human blood^6,7^**(Fig. 1b)**. In mice, ADH5 limits the toxicity of FA by converting it into the less toxic formate. To address whether cancer cells also rely on ADH5 activity to prevent FA toxicity, we inactivated the *ADH5* gene in HCT116 human colorectal carcinoma cells by CRISPR/Cas9 (**Extended Data Fig. 1a,b**). *ADH5*-deficient cells were not able to form tumor-spheroids in presence of FA, and they became sensitive to levels of FA near to those present in human blood **(Extended Data Fig. 1c,d)**. Moreover, ADH5 prevented the early apoptosis markers Annexin V, a blockage of the cell cycle at G2/M phase and sub-G1 DNA accumulation (**Fig. 1e,f and Extended Data Fig. 1c**), indicating that ADH5 limits FA-triggered cell death. Consistently, lymphoblastic leukemia Nalm6 cells lacking *ADH5* also presented strong sensitivity to blood FA levels, indicating that ADH5 protects unrelated human cancer cells from FA toxicity (**Extended Data Fig. 1d**).

### p53 orchestrates a FA response

Cell death can be a consequence of extensive damage to cellular components such as DNA, which might be detected by cell-fate regulators like p53^19^. Indeed, p53 has been shown to trigger a cellular response leading to acetaldehyde-mediated cell death in hematopoietic cells deficient in the Fanconi Anemia DNA crosslink repair pathway^10^. We set out to determine whether p53 could also orchestrate a cellular response to FA in HCT116 cells proficient for DNA repair leading to cell death. Surprisingly, the simultaneous inactivation of *P53* and *ADH5* only slightly suppressed the cytotoxicity of FA observed in HCT116 Δ*ADH5* cells (**Fig. 2a and Extended Data Fig. 1e,f**). In contrast, we found that inactivating *P53* significantly suppressed the severe colony-formation phenotype detected in Δ*ADH5* cells at FA concentrations as low as 12.5 μM but only mildly restored the formation of colonies at 25 μM FA (**Fig. 2b**), suggesting that FA can trigger cell death by both p53-dependent and independent pathways. HCT116 cells are proficient for the Fanconi Anemia DNA crosslink repair pathway, which might limit lethal FA genotoxicity. We therefore interrogated whether DNA damage was leading to a p53 response and to the accumulation of the double-strand break marker g-H2AX. We detected p53 phosphorylation, indicative of the activation of p53, in Δ*ADH5* but not in Wild type (WT) cells (**Fig. 2c,d and Extended Data Fig. 1e**), which correlated with cell cycle blockage at G2/M phase (**Fig. 1f**). However, we could not detect a significant induction of g-H2AX by blood-FA levels neither in WT nor in Δ*ADH5* cells (**Fig. 2c,d**). In contrast to FA treatment, exposure to the DNA-damaging drugs cisplatin, hydroxyurea (HU) or mitomycin C (MMC) resulted in a profound induction of those DNA-damage markers (**Fig. 2c,d**). To confirm that a 48-h exposure to micromolar levels of FA is not lethally genotoxic for cells proficient in DNA repair, we addressed genome instability by direct visualization of single chromosome damage (**Fig. 2e,f**). Indeed, we found that most of the metaphases in WT as well as in Δ*ADH5* cells were normal and only few of them presented chromosome damage. In stark contrast, severe chromosome damage was evident upon treatment with the DNA crosslinking agent mitomycin C, thus suggesting that when DNA repair is functional, FA might be causing cell death by damaging other cellular components than DNA.

**Fig. 2.**
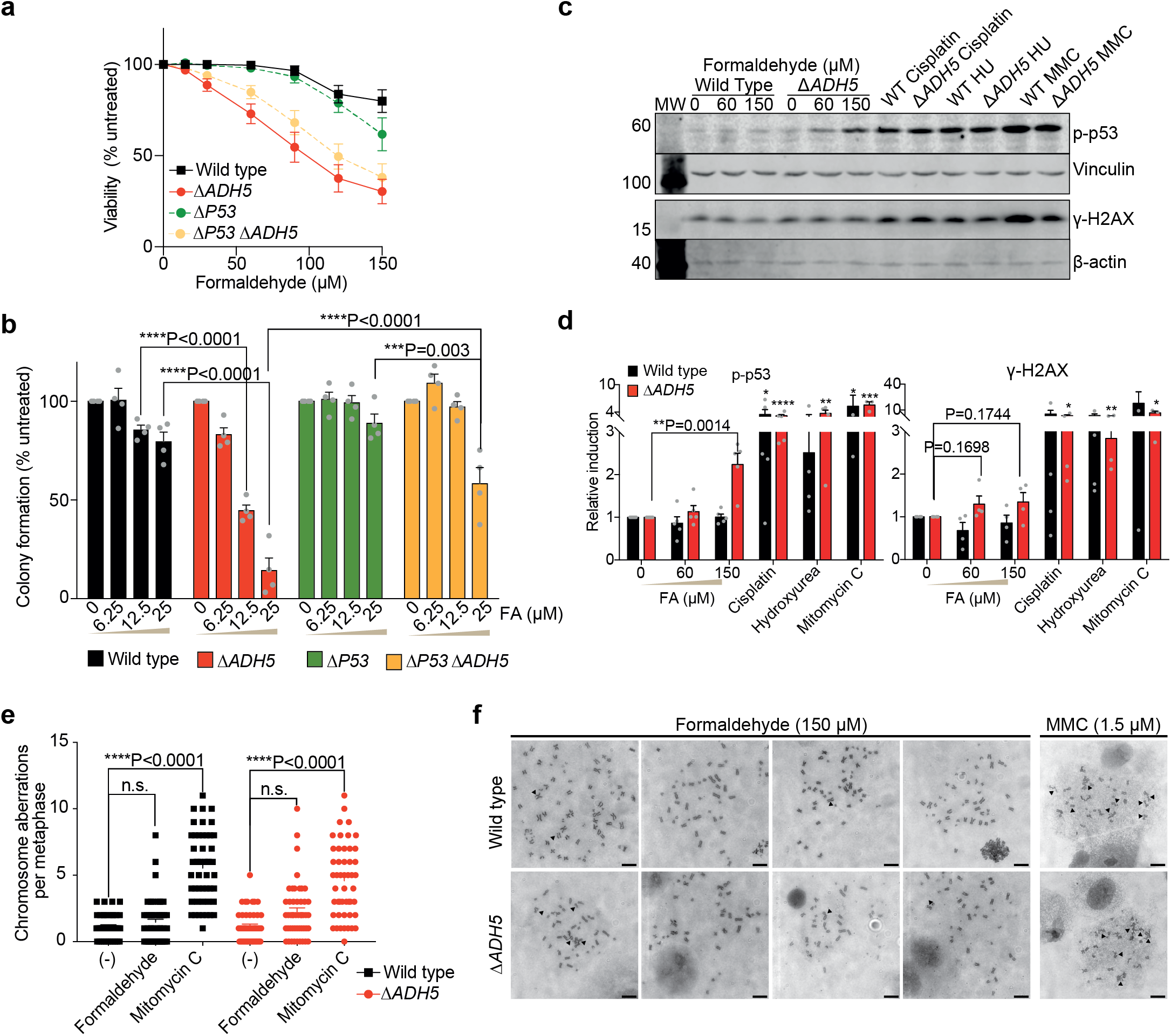
Formaldehyde triggers a p53 response in absence of *ADH5*. **a.** Resazurin-based viability assay expressed as % of fluorescence relative to untreated cells in Wild Type (WT), Δ*ADH5*, Δ*P53* and Δ*P53* Δ*ADH5* cells exposed to increasing concentrations of formaldehyde (FA) (mean ± s.e.m., n=8). **b.** Colony survival assay prepared seeding 600 cells in 6-well plates in presence of the indicated concentration of formaldehyde (FA). Colonies were stained and quantified after 10 days (mean ± s.e.m., n=4, one-way ANOVA using Tukey multiple comparison test). **c.** Western blot showing the induction of p-p53 and g-H2AX after 48 h of exposure to the indicated concentrations of FA or 24 h of exposure to the genotoxic compounds cisplatin (4 μM), hydroxyurea (HU, 1 mM) and mitomycin C (MMC, 1.5 μM). **d.** Quantitation of p-p53 and g-H2AX western blots using ImageJ (p-p53: FA 0, 60, 150 and cisplatin n=5; HU n=4, MMC n=3. g-H2AX: FA 0, 60, 150, cisplatin and HU n=4; MMC n=3; mean ± s.e.m., unpaired t-test comparing against the same cell line untreated) **e.** Quantitation of metaphase scoring (mean ± s.e.m., n=49, one-way ANOVA, Tuckey’s multiple comparison test) denoting the induction of chromosome damage by MMC (1.5 μM) but not by FA (150 μM). **f.** Representative images of Wild Type and Δ*ADH5* cells exposed to 150 μM FA and to 1.5 μM MMC (scale bar 1 μm).

### Oxidative stress underlies FA cytotoxicity

With the aim of discovering physiologically relevant cellular targets of FA, we reasoned that the strong avidity of the FA-carbonyl group toward electron-rich thiol groups might affect the antioxidant GSH. Indeed, the reaction between FA and the thiol group in GSH would block the redox capability of GSH, impairing its redox function. Moreover, the abundance of GSH (1-10 mM) might favor the spontaneous reaction between GSH and FA, which, if not limited, could diminish cellular GSH levels leading to oxidative stress. We therefore measured the cellular oxidative status by quantifying the oxidation of the probe 2’,7’-dichlorodihydrofluorescein diacetate (H2DCFDA). Interestingly, FA induced a significant oxidation of H2DCFDA in Δ*ADH5* cells (**Fig. 3a,b**). This oxidation level was comparable to that observed when exposing cells to the GSH-synthesis inhibitor L-buthionine-sulfoximine (L-BSO), and could be reverted by expressing ADH5 *in trans*. In order to test this more thoroughly, we incorporated the genetically-encoded cytosolic ROS sensor roGFP^20^.

**Fig. 3.**
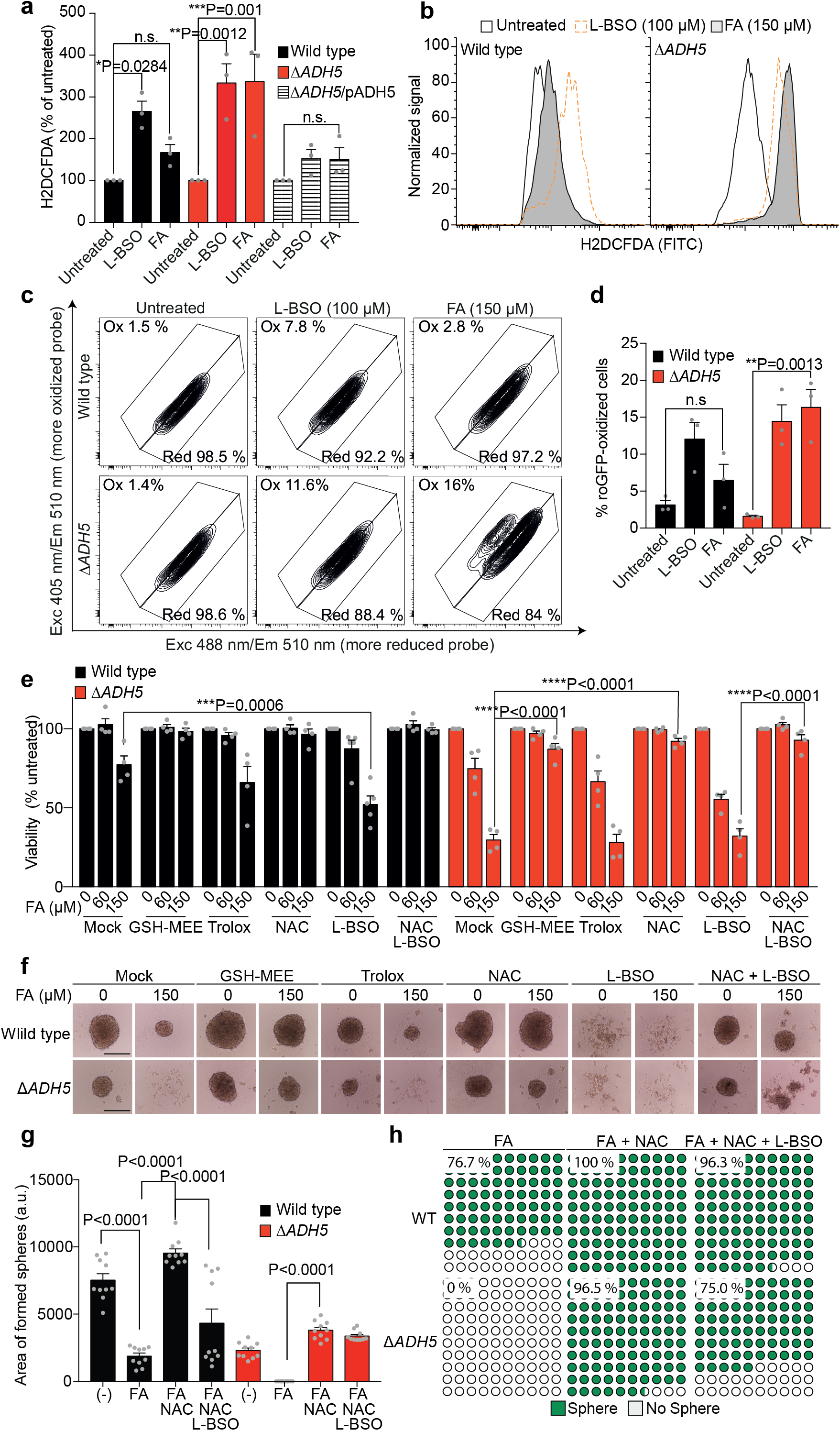
Oxidative stress underlies formaldehyde cytotoxicity. **a.** Oxidative stress determination by 2’,7’-dichlorodihydrofluorescein diacetate (HFDCDA) in Wild type, Δ*ADH5*, and complemented Δ*ADH5* (Δ*ADH5*/pADH5) HCT116 cells upon 48 h exposure to formaldehyde (FA). Data is represented as the % of fluorescence detected in the untreated samples from the same cell line (mean ± s.e.m., n=3, one-way ANOVA, Tuckey’s multiple comparison test). **b.** Representative plots of oxidative stress determination as described in a. **c.** Flow cytometry representative plots obtained from Wild type and Δ*ADH5* cells harboring the cytoplasmic-roGFP reporter. Cells were excited at λ = 405 or λ = 488 nm and emission recorded at λ = 510 nm. **d.** Quantitation of experiments shown in d (mean ± s.e.m., n=3, one-way ANOVA, Tuckey’s multiple comparison test). **e.** Resazurin-based viability at 0, 60 and 150 μM FA denoting the rescue of cell viability by N-acetylcysteine (NAC, 500 μM), Trolox (1 mM) and glutathione monoethyl ester (GSH-MEE, 1 mM). L-buthionine-sulfoximine (L-BSO) was used at 100 μM. The data represent the mean ± s.e.m. of 4 experiments done in triplicate (one-way ANOVA, Tuckey’s multiple comparison test). **f.** Representative images of the 3D-tumour spheroid formation phenotype in presence of the indicated FA concentration and the combination of the antioxidants described in e (scale bar 0.5 mm). **g.** Quantitation of 3D-sphere area from 10 formed spheres at day 5 after seeding (mean ± s.e.m., one-way ANOVA, Tuckey’s multiple comparison test). **f.** Quantitation of sphere formation phenotype at day 5 after seeding 2000 cell/well of WT or Δ*ADH5* cells in presence of FA (150 μM); FA and NAC (500 μM); or FA, NAC and L-BSO (100 μM). The plots correspond to a part of the whole representation (WT + FA, n=30; Δ*ADH5* + FA, n=28; WT + FA + NAC, n=29; Δ*ADH5* + FA + NAC, n=29; WT + FA + NAC + L-BSO, n=27; Δ*ADH5* + FA + NAC + L-BSO, n=28).

Exposure to FA induced a population of cells in which the sensor is oxidized in the absence of *ADH5*. These results indicate that FA detoxification is necessary to prevent oxidative stress (**Fig. 3c,d**).

To address the causal contribution of FA-induced oxidative stress to cell death, we set out to test whether cell toxicity could be rescued by the antioxidants N-acetylcysteine (NAC), glutathione monoethyl ester (GSH-MEE) or Trolox (water-soluble vitamin E). The death phenotype and the 3D-sphere formation defect could be almost fully reverted by incubating with GSH-MEE or NAC, indicating that an increase in free-thiols can prevent FA cytotoxicity (**Fig. 3e,f**). Remarkably, GSH-MEE and NAC led to an overgrowth of WT 3Dspheres (**Fig. 3f,g**). In contrast, Trolox, a non-thiol antioxidant, was unable to limit FA toxicity, suggesting that oxidative stress *per se* is not sufficient to poison Δ*ADH5* cells (**Fig. 3e,f**). NAC can work by directly scavenging free FA or by boosting endogenous GSH^21^. To further interrogate the suppressive effect observed with this thiol-rich antioxidant, we combined L-BSO and NAC. Remarkably, NAC could still rescue the toxicity caused by FA even when GSH synthesis was inhibited by L-BSO (**Fig. 3e**). However, blocking GSH synthesis limited the overgrowth phenotype observed in 3D-spheres exposed to NAC (**Fig. 3e,g**). Moreover, GSH synthesis inhibition reduced the NAC-rescue of 3D-sphere formation in Δ*ADH5* cells exposed to FA from 96.5 % to 75 % (**Fig. 3h**). Altogether, these observations indicate that ADH5 limits oxidative stress induction by FA and that supplying GSH can prevent FA toxicity.

### GSH biosynthesis limits FA toxicity

Exogenous GSH precursors can prevent FA toxicity; we therefore predicted that limiting endogenous GSH should increase FA toxicity even in presence of ADH5. First, we selected concentrations of the GSH synthesis inhibitor L-BSO that were not cytotoxic to the human cancer cells HCT116 and Nalm6 (**Extended Data Fig. 2a**). The viability of WT HCT116 and Nalm6 cells in presence of FA was significantly reduced in presence of L-BSO, indicating that GSH synthesis contributes to cellular FA tolerance (**Fig. 3e,4a**). Surprisingly, a non-cytotoxic L-BSO concentration affected the formation of 3D-spheres in both WT and Δ*ADH5* HCT116 cells even in absence of exogenous FA (**Fig. 4b**). In Nalm6 cells, which grow in suspension, the treatment with L-BSO increased the sensitivity of Δ*ADH5* cells to FA (**Fig. 4a**), suggesting that GSH biosynthesis and ADH5 independently contribute to prevent FA toxicity in this lymphoblastic human cancer cell. Although L-BSO is neither cytotoxic to HCT116 nor Nalm6 cells at the concentrations used in our experiments (**Extended Data Fig. 2a**), it is still a pharmacological avenue that might have off-target effects. We therefore set out to genetically inactivate GSH biosynthesis (*GCLM*) by CRISPR/Cas9 in HCT116 cells (**Extended Data Fig. 2b,c,d**). Concordantly with the pharmacological experiments, *GCLM* deficiency reduced cellular tolerance to FA (**Fig. 4c**). The simultaneous inactivation of *ADH5* and *GCLM* did not further affect viability (**Fig. 4c**). This result indicates that for cell viability, ADH5 is the dominant factor in protecting HCT116 cells against FA. The 3Dsphere formation phenotype was affected by the sole inactivation of *GLCM* (**Fig. 4d**), concordantly with the results observed using the GSH-synthesis inhibitor L-BSO (**Fig. 4b**). In contrast, the formation of colonies was further impaired in Δ*GCLM* Δ*ADH5* cells compared to single knockout counterparts, thus revealing an independent contribution of GSH biosynthesis and ADH5 to this phenotype (**Fig. 4e,f**). The disparity observed in viability and colony survival assays might indicate that in absence of both *ADH5* and *GCLM* some phenotypes such as cell-cell interaction might be affected without necessarily impairing cell viability.

**Fig. 4.**
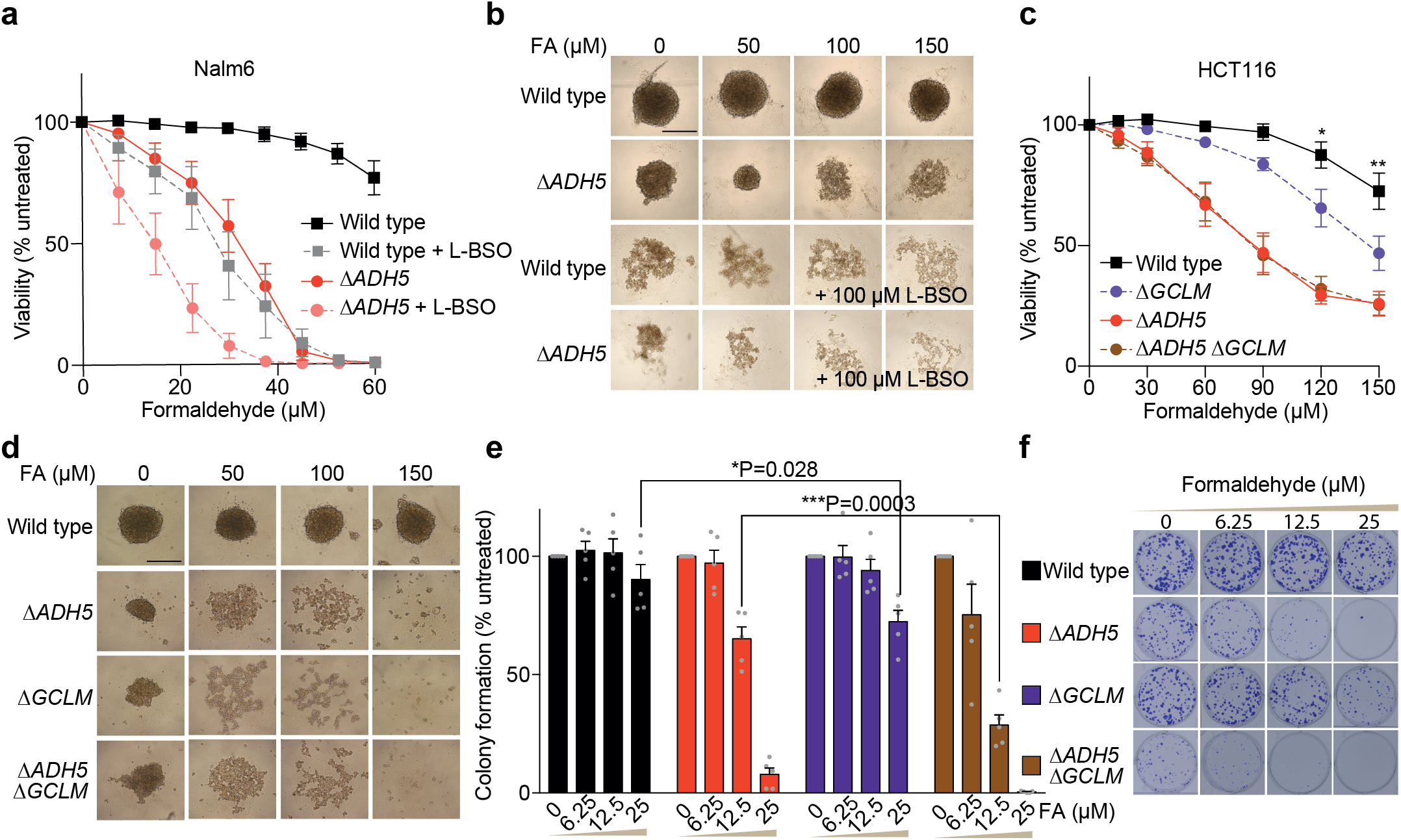
GSH biosynthesis limits formaldehyde toxicity. **a.** Resazurin-based viability assay in Wild type (WT) and Δ*ADH5* Nalm6 cells in presence of different concentrations of formaldehyde (FA) with or without 50 μM L-buthionine-sulfoximine (L-BSO) (mean ± s.e.m., n=6). **b.** HCT116 3D-spheroid formation in presence of 100 μM L-BSO and the indicated concentrations of FA. Pictures were taken 5 days after seeding cells on agarose-coated plates (scale bar 0.5 mm). **c.** Resazurin-based viability assay performed with Wild type, Δ*ADH5*, Δ*GCLM* and Δ*ADH5* Δ*GCLM* cells in response to increasing concentrations of FA (mean ± s.e.m., n=5, asterisks represent the statistical significance according to one-way ANOVA for multiple comparison using a Tukey-corrected test between Wild type and Δ*GCLM*). **d.** Representative images of HCT116 3D-spheroid formation for the same cell lines described in (C). Pictures were taken at day 5 after seeding (scale bar 0.5 mm). **e.** Colony survival assay prepared seeding 600 of WT, Δ*ADH5*, Δ*GCLM* and Δ*GCLM* Δ*ADH5* cells in 6-well plates in presence of the indicated concentration of FA (mean ± s.e.m., n=5, one-way ANOVA using a Tukey’s multiple comparison test). **f.** Representative images of the colony survival assay quantified in e.

### Endogenous FA reacts with GSH yielding HSMGSH

GSH and FA metabolisms are linked as FA spontaneously reacts with GSH yielding HSMGSH (Fig. 5A), a substrate of ADH5. We hypothesized that this reaction might occur *in vivo* affecting the endogenous level of GSH as well as limiting the reactivity of free FA. By *in house* synthesis and reaction monitoring using ultraperformance liquid chromatography coupled to high resolution mass spectrometry (UPLC-HRMS), we first confirmed that GSH and FA react *in vitro* yielding HSMGSH, which was subsequently used as chemical standard (**Extended Data Fig. 3**). Should cellular metabolism generate endogenous FA, we might be able to detect the formation of HSMGSH. By UPLC-HRMS, we were able to detect this compound together with GSH and GSSG in cell extracts (**Fig. 5b,c and Extended Data Fig. 4a-d**). The continuous generation of FA from cellular metabolism might need a constant recovery of reduced GSH from HSMGSH formation to sustain endogenous GSH. Indeed, cells lacking *ADH5* presented significantly lower levels of GSH compared to WT cells (**Fig. 5d**). This decrease is in line with the accumulation of HSMGSH relative to GSH (**Fig. 5e**). However, the net amount of total GSH and HSMGSH was lower in Δ*ADH5* cells, thus we cannot rule out the participation of efflux mechanism(s) pumping out HSMGSH when this product accumulates (**Fig. 5e,f**). To confirm that Δ*ADH5* cells present lower levels of GSH, we interrogated GSH by using an indirect fluorescent reagent. According to this assay, cells lacking *ADH5* contained 17.9 % less reduced GSH than the WT counterparts, corroborating that *in vivo* ADH5 significantly contributes to cellular GSH (**Fig. 5g**). The genetic inactivation of the regulatory component in the rate-limiting step of GSH biosynthesis (*GCLM*) or the treatment with L-BSO further depleted endogenous GSH in both Δ*ADH5* and WT cells, denoting that the mechanism by which ADH5 contributes to GSH homeostasis is downstream GSH synthesis (**Fig. 5g**).

**Fig. 5.**
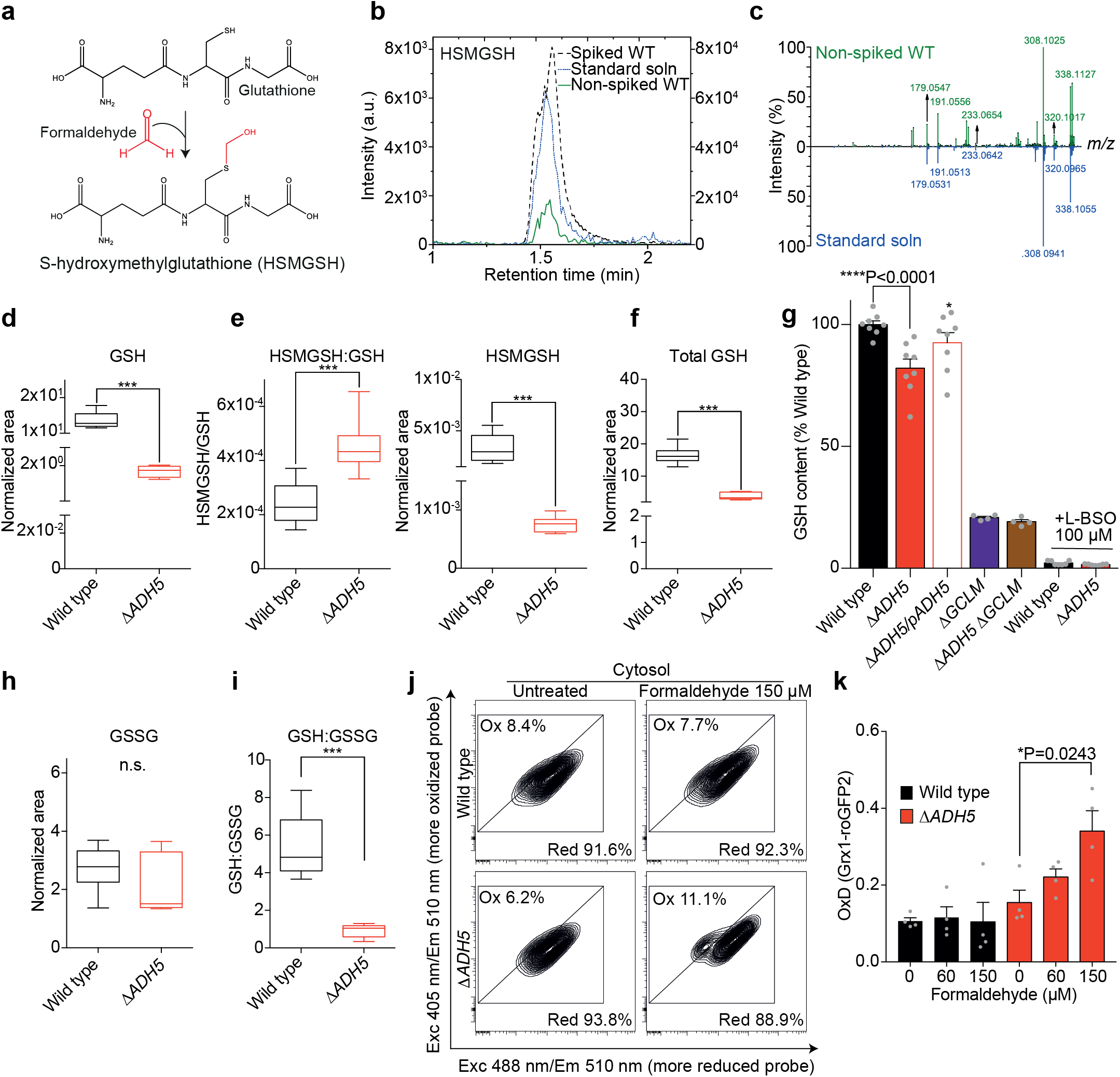
Endogenous formaldehyde reacts with GSH altering the GSH:GSSG ratio. **a.** Scheme showing the spontaneous reaction between formaldehyde (FA) and glutathione (GSH) yielding S-hydroxymethylglutathione (HSMGSH). **b.** Extracted ion chromatograms for [HSMGSH + H]+ ion at m/z 338.1022 ± 0.0500 generated from a non-spiked Wild type (WT) sample (green, left axis), a 20 μM spiked WT sample (black dotted line, right axis), and a 20 μM HSMGSH standard solution (blue, left axis). **c.** Product ion mass spectra of [HSMGSH + H]^+^ precursor ion for a WT sample (green), and for a 20 μM HSMGSH standard solution (blue), using a collision cell voltage of 10 V. **d.** Box and whiskers plot for GSH content in WT and Δ*ADH5* cells calculated as normalized chromatographic peak areas relative to the number of viable cells (n=9, Mann-Whitney test, *** P<0.0001). The box and whiskers plots are represented by a line in the box corresponding to the median; the edges are the 25th and 75th percentiles and the whiskers extend to the most extreme values in data. **e.** Left: Box and whiskers plot for HSMGSH content relative to GSH in WT and Δ*ADH5* cells (n=9, Mann-Whitney test). Right: Net HSMGSH content in WT and Δ*ADH5* cells calculated as normalized peak areas relative to the number of viable cells (n=9, Mann-Whitney test). **f.** Box and whiskers plot for total GSH (GSH disulfide (GSSG) plus GSH) content in WT and Δ*ADH5* cells calculated as normalized peak areas relative to the number of viable cells (n=9, Mann-Whitney test). **g.** Bar plots for GSH content in WT (n=17), Δ*ADH5* (n=15), Δ*ADH5*/pADH5 (n=8), Δ*GCLM* (n=4) and Δ*ADH5* Δ*GCLM* (n=4) cells, and in WT (n=7) and Δ*ADH5* (n=7) cells exposed to 100 μM L-BSO for 48 h. Every dot is the percentage of fluorescence intensity in a single well relative to the average fluorescence of WT samples run the same day and corrected for viability determined using resazurin (mean ± s.e.m., unpaired t-test). **h.** Box and whiskers plot for GSSG content in WT and Δ*ADH5* cells calculated relative to the number of viable cells (n=9, Mann-Whitney test). **i.** Box and whiskers plot for GSH:GSSG ratio in WT and Δ*ADH5* cells (n=9, Mann-Whitney test). **j.** Flow cytometry representative plots from WT and Δ*ADH5* cells harboring the cytosolic Grx1-roGFP2 reporter. Data was recorded 48 h post-FA exposure. **k.** Quantitation of oxidized Grx1-GFP2 (OxD (Grx1-roGFP2)) sensor from plots depicted in (J) (mean ± s.e.m., n=4, unpaired t-test).

### HSMGSH metabolism prevents GSH:GSSG imbalance

In the cytosol, GSH and GSSG levels have been reported to be around 10 mM and 200 nM, respectively, determining a cytosolic GSH redox potential (E_GSH_) of −320 mV^13^. Despite the high level of reduced GSH, a small change in the ratio between the reduced and the oxidized GSH form (GSH:GSSG) can substantially affect E_GSH_, thus impairing cellular redox balance^22^. We reasoned that blocking the GSH supply through ADH5 would affect the GSH:GSSG ratio, which might consequently lead to oxidative stress (**Fig. 3a,d**). We therefore measured relative levels of GSSG in both WT and Δ*ADH5* cells (**Fig. 5h**) and calculated the GSH:GSSG ratio from UPLC-HMRS data (**Fig. 5i**), observing a 5.9-fold reduction in Δ*ADH5* compared to WT cells (**Fig. 5i**). To interrogate the role of *ADH5* in maintaining the GSH:GSSG ratio upon FA stress, we incorporated the cytoplasmic version of the reporter Grx1-roGFP2^23^ in HCT116 WT and Δ*ADH5* cells. This ratiometric reporter (λ∈m = 510 nm) contains two cysteines that can form a reversible disulfide bond that is in equilibrium with the endogenous GSH:GSSG couple. In a more oxidant environment, the ratio between GSH and GSSG will decrease leading to a more oxidized Grx1-roGFP2 sensor. The fraction of the oxidized sensor (OxD Grx1-roGFP2) can be calculated from the ratio between the Grx1-roGFP2 emission at λ = 510 nm when it is excited at λ = 405 and λ = 488 nm (R405/488)^23^. We found that ADH5 prevented the FA-dependent oxidation of Grx1-roGFP2 in the cytosol (**Fig. 5j**), concordantly with the detection of H2CDFDA and roGFP oxidation (**Fig. 3a-d**). In summary, these results show that HSMGSH metabolization by ADH5 can prevent cytoplasmic GSH:GSSG imbalance by supplying cellular GSH.

### The role of ADH5 is conserved

In order to interrogate the relevance of GSH metabolism and ADH5 beyond human cancer cells, we explored the presence of genes coding for ADH5-like proteins in the metazoan model *Caenorhabditis elegans* (**Extended Data Fig. 5a**). In the nematode, the uncharacterized gene H24K24.3 codes for the ortholog of the human ADH5 enzyme. Transgenic expression of ADH-5 fused with GFP under the control of the endogenous *adh-5* promoter (Ex[*p_adh-5_*ADH-5::GFP; *p_myo-2_*tdTomato]) presented a ubiquitous cytoplasmic expression in larvae and in the adult nematode (**Fig. 6a**). To assess whether H24K24.3 participates in the prevention of FA toxicity in worms, we generated a null mutant via CRISPR/Cas9 by introducing multiple stop codons in all three reading frames^24^. Animals lacking H24K24.3 showed an extreme hypersensitivity to FA (**Fig. 6b,c**), affecting the survival throughout development (**Extended Data Fig. 5b**), overall indicating that H24K24.3 is the ortholog of *ADH5* in *C. elegans*. We thus refer to H24K24.3 from now on as *adh-5*. While *adh-5(sbj21)* mutant *C. elegans* larvae did not survive FA exposure, a pre-treatment with only 10 μM NAC significantly restored survival of *adh-5* mutants and also allowed animals to develop into adulthood, assessed 72 h post FA treatment (**Fig. 6b-d and Extended Data Fig. 5b,c**). Conversely, treatment of L1 larvae with a sublethal FA concentration and simultaneous exposure to the prooxidant paraquat (PQ), which generates ROS in *C. elegans*^25^, severely affected the development of L1 *adh-5* larvae (**Fig. 6e,f**). These results indicate that providing an antioxidant can reduce FA toxicity, while additional oxidative damage increases FA stress in nematodes, strongly supporting our model of oxidative GSH imbalance as a FA-cytotoxic effect.

**Fig. 6.**
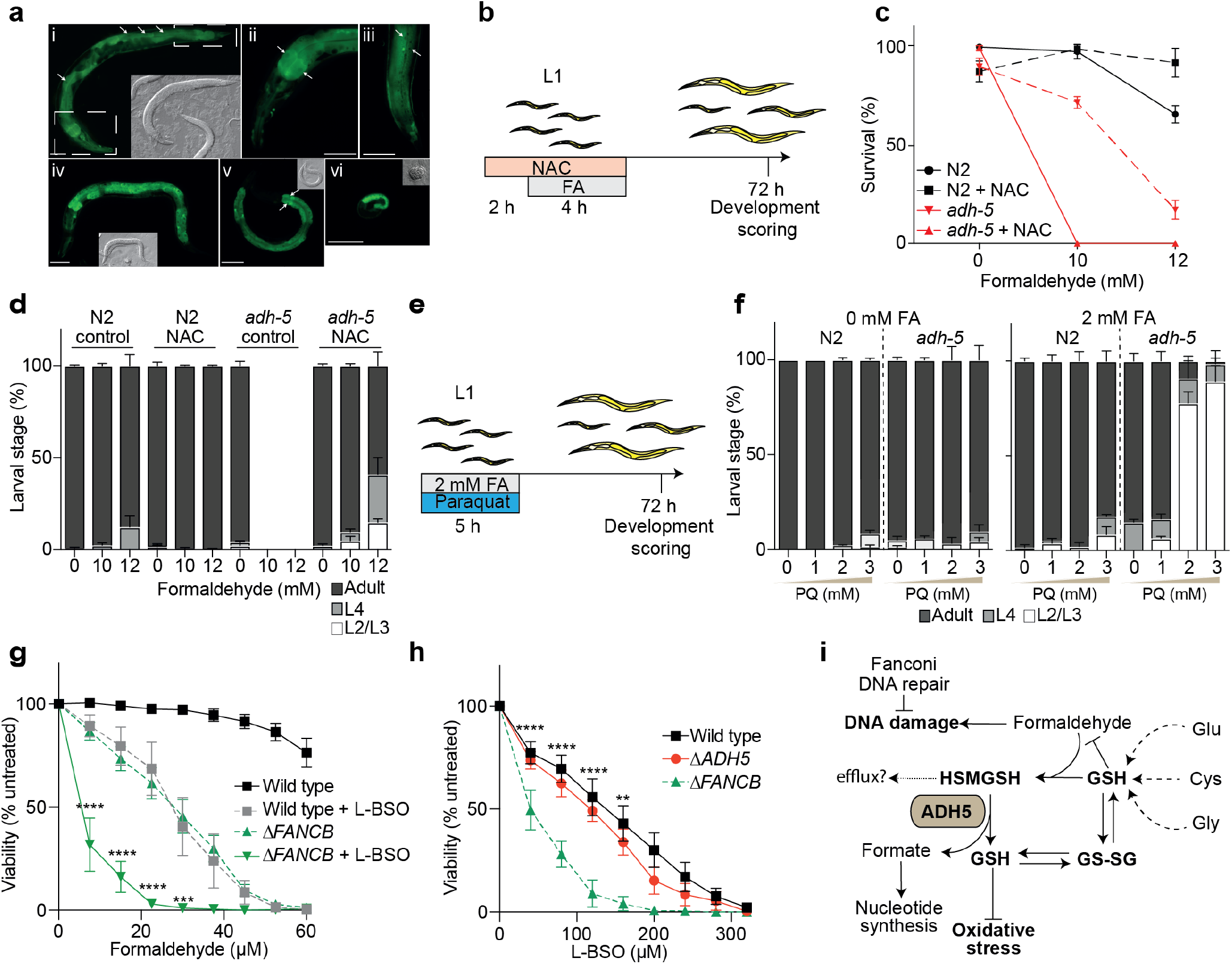
The *ADH5* ortholog in *C. elegans* and GSH synthesis in Fanconi Anemia-deficient cells. **a.** Various developmental stages of a *C. elegans* transgenic line expressing *Fx*[p_*adh-5*_ADH-5::GFP; p_*myo-2*_tdTomato]. White arrows point to nuclei. Scale bars represent 50 μm. Gray inlays show the corresponding DIC image. (i) Adult stage boxes refer to ii and iii. (ii) Head area with focus on the intestine of the adult depictedin i, recorded with 40x magnification. (iii) Tail area with focus on the cuticle of the animal shown in i, recorded with 40x magnification. (iv) L4 stage. (v) L3 stage. (vi) Late embryo. **b.** Scheme depicting the protocol used to treat *C. elegans* with formaldehyde (FA) and N-acetylcysteine (NAC). **c.** Survival of L1-staged Wild type (N2) and *adh-5* mutant upon exposure to the indicated FA concentrations and 10 μM NAC measured directly after treatment (mean ± S.D., n=3). **d.** Development profile of surviving animals 72 h after FA exposure (mean ± S.D., n=3). **e.** Scheme depicting the protocol used to treat *C. elegans* with FA and paraquat (PQ). **f.** Development profile of surviving animals 72 h upon FA and PQ exposure (mean ± S.D., n=3). **g.** Resazurin-based viability assay for Nalm6 cells exposed to increasing concentrations of FA and 50 μM L-buthionine-sulfoximine (L-BSO), denoting the protection against FA by glutathione (GSH) biosynthesis (mean ± s.e.m., n=6, asterisks represent one-way ANOVA for multiple comparison using a Tukey-corrected test between Δ*FANCB* and Δ*FANCB* + L-BSO 50 μM). **h.** Resazurin-based viability assay for Nalm6 cells exposed to increasing concentrations of L-BSO (mean ± s.e.m., n=6, asterisks represent the statistical significance according to one-way ANOVA for multiple comparison using a Tukey-corrected test between Wild type (WT) and Δ*FANCB* cells). **i.** Model for FA metabolism and the crosstalk with GSH metabolism highlighting the formation of HSMGSH adducts and the metabolization through ADH5. GSH supply by ADH5 limits oxidative stress and sustains the balance between GSH and GSH disulfide (GSSG).

Finally, we reasoned that the reaction between FA and GSH forming HSMGSH molecules and their metabolization by ADH5 might limit free FA. It has been shown that cells lacking the interstrand-crosslinking (ICL)-DNA repair pathway Fanconi Anemia are very sensitive to FA^9–11^. We therefore predict that GSH will be required to prevent FA toxicity in Fanconi Anemia. To assess our hypothesis, we exposed Nalm6 cells deficient in *FANCB*, a Fanconi Anemia DNA crosslink repair gene, to FA in presence of L-BSO. As predicted, Δ*FANCB* cells were sensitive to FA and this phenotype was largely exacerbated by blocking GSH synthesis (**Fig. 6g**). Interestingly, cells lacking *FANCB* were significantly sensitive to GSH inhibition even in absence of FA, which might be a consequence of an increase in endogenous free FA, overall suggesting that GSH supply might be fundamental for Fanconi Anemia patients (**Fig. 6h**).

## Discussion

In this work we reveal that FA can cause cytotoxicity by triggering oxidative stress (**Fig. 3a-d**), explaining earlier observations of oxidative damage in tissues and cells exposed to FA^26,27^. We determined that FA reacts with GSH, affecting the GSH:GSSG ratio and the cellular redox balance. We describe a conserved mechanism to salvage GSH from FA-GSH covalent products (HSMGSH) limiting FA cytotoxicity not only in cancer cells but also in *C. elegans*. This pathway is centered on the enzyme ADH5 and it is downstream of the *de novo* GSH synthesis pathway (**Fig. 6i**).

At physiological FA concentrations as they occur in human blood, ADH5 is essential for cellular growth and viability (**Fig. 1c,d, 2c,d**). In Δ*ADH5* cells FA treatment triggers p53 activation that accounts for some aspects of the FA response such as proliferation arrest at non-cytotoxic FA concentrations (**Fig. 2b**), while p53 is dispensable for the decline in viability (**Fig. 2c**) suggesting other cell-fate regulators may trigger FA-induced cell death. It is likely that FA-induced DNA damage upregulates a p53 response that blocks cell cycle at G2/M phase, thus impairing the formation of colonies. However, in presence of functional DNA repair mechanisms, DNA damage would be alleviated before it reaches the threshold required for triggering p53-dependent apoptosis. On the other hand, the FA-induced metabolic disruption might lead to p53-independent cell death. Further research should reveal the identity of the p53-independent mechanisms that respond to FA.

The detection of HSMGSH in cells not exposed to exogenous FA indicates that cellular metabolism produces sufficient FA to react with GSH yielding HSMGSH (**Fig. 5b,c**). ADH5 restores GSH by metabolizing HSMGSH and thus maintaining the cellular GSH balance to limit oxidative stress. Several factors -in addition to GSH-have been implicated in the cellular protection against oxidative stress, most of them being under the control of the master regulator NRF2^28^. Upon detecting oxidative stress, the NRF2 partner KEAP1 no longer ubiquitin-labels NRF2 for degradation, resulting in NRF2 stabilization and activation of NRF2-response genes. Indeed, inactivating *Keap1* has been shown to rescue a phenotype of diet-induced steatohepatitis reported in *Adh5*^-/-^ mice^29^. NRF2 is a tumor suppressor gene that also controls stem cell fate and the crosstalk between NRF2 and HSMGSH metabolism might have significant consequences beyond cancer.

ADH5 can also metabolize S-nitrosoglutathione (GSNO) producing ammonia and GSH^12^. This enzymatic activity gave origin to the alternative name GSNOR and it has prompted the development of pharmacological inhibitors that might be used for modulating nitric oxide homeostasis in inflammatory diseases^30^. Remarkably, GSH is the common product of the enzymatic activity of ADH5 using either HSMGSH or GSNO as substrates. Thus, blocking ADH5 might trigger adverse effects such as GSH redox imbalance and increased toxic endogenous FA. On the other hand, GSH biosynthesis has been explored as a therapeutic target to overcome resistance to cancer combinatorial therapies. However, cancer cells can compensate GSH depletion by inducing the thioredoxin (TXN) pathway, which helps to maintain cellular antioxidant capacity^31^, and by maintaining protein homeostasis through deubiquitinating enzymes (DUB)^32^. Since FA was shown to induce a proteotoxic stress response^33^, thus ADH5 inhibition might improve the efficacy of DUB inhibitors and GSH depletors in cancer therapy.

Our findings may have wide implications for the human disease Fanconi Anemia. Metabolic ROS were shown to induce DNA damage in hematopoietic stem cells (HSCs) when they start cycling to exit quiescence, which impairs blood production in *Fanca*-/- mice^34^. It is known that oxygen can exacerbate chromosome aberrations in lymphocytes from Fanconi Anemia patients^35^. Moreover, the GSH precursor NAC has been shown to improve genome stability in these lymphocytes^36^. It is likely that an increase in ROS as consequence of oxygen exposure would affect GSH pool, thus indirectly leading to accumulation of FA and genome instability. A combined therapy using a FA sponge such as metformin^37^ and GSH-precursors might succeed in benefiting Fanconi Anemia patients. Furthermore, a diet rich in GSH-precursors might delay cancer onset in healthy *BRCA2*-mutation carriers by limiting FA toxicity, overall highlighting the broad reach of the findings reported here.

## Methods

### Experimental Model and Subject Details

#### Cells and animals

HCT116 cells were maintained in Dulbecco’s Modifies Eagle’s Medium (DMEM) high glucose (Thermo Scientific, #12100061), supplemented with 1 % Penicillin/Streptomycin and 10 % FBS (Natocor)^38^. Nalm6 cells were maintained in Roswell Park Memorial Institute 1640 medium (RPMI) (Thermo Scientific, #31800105)^3^ containing 10 % FBS, 1 % Penicillin/Streptomycin and 50 μM ß-mercaptoethanol. All the cell lines were regularly tested for mycoplasma infection.

Housing and handling of mice were performed in agreement with animal protection guidelines of the district president of Cologne. All procedures were approved and authorized by the LANUV with identification number 84-02.04.2015.A484. ‘Role of ageing-associated DNA damage in energy homeostasis-regulating neurons. Mice were maintained in individually ventilated cages (IVCs) on autoclaved bedding and food and sterile-filtered water in a barrier facility at the University of Cologne and the MPI for Metabolism Research. Mice were subjected to a constant 12-h day–night cycle and a constant room temperature of 22°C. Starting from 2 months of age, mice were fed a control diet (CD) consisting of 67 kJ % carbohydrates, 20 kJ % proteins and 13 kJ % fat (Sniff). Mice had ad libitum excess to food and water. The NPY-GFP mice were backcrossed for at least two generations to the C57BL/6N background^39^.

*Caenorhabditis elegans* was maintained using standard methods^40^. N2, Bristol *C. elegans* wild isolate was obtained from *Caenorhabditis* Genetics Center (CGC), Minneapolis, MN, USA.

#### CRISPR/Cas9 generation of Δ*ADH5* and Δ*GCLM* cell lines

HCT116 Δ*ADH5*, Δ*GCLM*, Δ*P53* Δ*ADH5* and Δ*GCLM* Δ*ADH5* cell lines were generated by targeting exon 3 of *ADH5* (sgRNA: TGCTGGAATTGTGAAAGTGTT) and exon 1 of *GCLM* (sgRNA: ACGGGGAACCTGCTGAACTG) in the corresponding parental cell lines. Briefly, sgRNAs were cloned into the pX458 vector (Addgene, #48138) and transfected using lipofectamine 3000 (Thermo Scientific, L3000015). GFP-expressing cells were sorted and clonally diluted in 96-well plates. After 20 days, cells were expanded, and inactivation of the desire gene confirmed by western blot using GCLM (Atlas antibodies, #HPA023696) or ADH5 antibodies^9^. The mutations generated by Cas9 at the target exons were obtained by preparing genomic DNA from the selected clones and amplifying the exons with the primers: Fwd (hA5-ck1): 5’-TCTTGTATCTGTACCTCTGA-3’; Rv (hA5-ck1rv): 5’-CCTTCAGCTTAGTAACTC-3’ for *ADH5*, and Fwd (hGCLM-ck_834Fw): 5’-GAAGCACTTTCTCGGCTACG-3’; Rv (hGCLM-834_Rv): 5’-TCCTTTACCTGGACAGGGTG-3’ for *GCLM*. PCR results were analyzed by gel electrophoresis, cloned and sequenced using universal M13 primers.

#### Generation of cells stably expressing ADH5

Cells carrying the ADH5-expressing plasmid pLox-ADH5-FLAG-CT-BSR^3^ were selected using 4 μg/ml Blasticidin (BSR). BSR-resistant cells were clonally diluted and ADH5-expression verified by western blot against FLAG epitope.Cells carrying the ADH5-expressing plasmid pLox-ADH5-FLAG-CT-BSR^3^ were selected using 4 μg/ml Blasticidin (BSR). BSR-resistant cells were clonally diluted and ADH5-expression verified by western blot against FLAG epitope (Abcam, #ab49763).

#### Generation of *C. elegans* lines

The ADH-5::GFP reporter line *Ex*[p_*adh-5*_ADH-5::GFP; p_*myo-2*_tdTomato] was produced via co-injecting the clone (5736523864883943 G06, tagged gene: H24K24.3) of the TransgeneOme fosmid library^41^ together with the selection marker for tdTomato expression in the pharynx, using standard *C. elegans* microinjection^42^. For imaging, various stages of the transgenic animals were mounted on 5 % agar pads with polystyrene nanoparticles (Polysciences, 2.5 % by volume) as previously described^43^ and imaged at an AxioImager M.2 fluorescence microscope (Zeiss, Jena, Germany).

The *C. elegans* orthologue of human *adh-5* gene (H24K24.3) was knocked out using the CRISPR/Cas9 system as previously reported: The preassembled CRISPR/Cas9 ribonucleoprotein complexes and linear single stranded DNAs as repair templates were directly injected into the gonad of young adult hermaphrodites^44^. To generate *adh-5* null mutant, we utilized a universal STOP-IN cassette that contained an exogenous Cas9 target site, multiple stop codons in all three reading frames and the recognition site of the *NheI* restriction enzyme^24^. The *C. elegans adh-5* sgRNA with GGG protospacer adjacent motif was designed using Benchling (https://benchling.com/) and targeted exon 3 of the *adh-5* gene (5’-CTTCATGTCCCAAGACGACA-3’). The *C. elegans* DNA repair oligo included a STOP-IN cassette and two short homology arms identical to the sequences flanking the Cas9 cleavage site (5’-GCCACACGGACGCCTACACCCTCGACGGACACGATCCGGAAGGTCTCTTCCCTGTGGGAAGTTTGTCCAGAGCAGAGGTGACTAAGTGATAAGCTAGCCGTCTTGGGACATGAAGGGTCTGGAATTGTCGAGA-3’). To facilitate screening, a co-conversion strategy with dominant phenotypic roller marker was used^45^. Microinjection was performed as previously described^42^ using the following injection mix: KCl (25 mM), Hepes pH 7.4 (7.5 mM), tracrRNA (200 ng/μl), *dpy-10* crRNA (150 ng/μl), *dpy-10* ssODN (13.75 ng/μl), *adh-5* sgRNA (300 ng/μl), *adh-5* ssODN (100 ng/μl), Cas9 (416 ng/μl, NEB, USA). F1 worms carrying roller phenotype were preselected and cloned 4-6 days after the injection. The F2 progeny was subsequently screened for the desired edit by PCR amplification using the *adh-5* forward primer (5’-CGATCCAAGTGGCTCCACCGAA-3’) and the *adh-5* reverse primer (5’-TTCCACATCCCAAAAGCGAAACC-3’). The presence of the STOP-IN cassette was verified via Sanger sequencing (Eurofins Genomics, Germany) with the *adh-5* sequencing primer (5’-CGATTAACCGACACCCTTGCTC-3’).

#### Survival and development assays in *C. elegans*

For the combined N-acetylcysteine (NAC, Sigma-Aldrich, #A7250) and formaldehyde (FA, Pierce, #28908) treatment worm stages were first synchronized via bleach-synchronization. Gravid adult animals and eggs were harvested from NGM plates with 5 ml M9 buffer (3 g KH_2_PO_4_, 6 g Na_2_HPO_4_, 5 g NaCl, in 1 l H_2_O; autoclaved and added 1 ml 1 M MgSO_4_) using a cell scraper and transferred to 15 ml tubes, before adding 1 ml bleach solution (5M NaOH and sodium hypochlorite in a 1:1 ratio). The tubes were then constantly vortexed for 5 min and centrifuged (Centrifuge 5810R, Eppendorf) at 2800 rpm for 1 min. After removing the supernatant, the worms were washed three times with 5 ml M9 medium by shaking the tubes and then centrifuging at 2800 rpm for 1 min. Finally, they were kept in 10 ml M9 medium overnight (16 h) under rotation at 35 rpm (Multiple-Axle-Rotating-Mixer RM10W-80V, CAT) to allow animals to hatch. Prior to FA treatment, L1-staged worms were centrifuged at 1300 rpm for 1 minute, and the volume was reduced to 1 ml M9 medium. The number of worms was determined under a stereoscope in a representative volume of 3 μl and a final concentration of approx. 50-100 worms per μl was adjusted. A solution was prepared by pelleting a saturated OP50 *E. coli* bacterial culture, which was first heat-inactivated (60 °C O/N), at 4000 rpm for 10 minutes, and then concentrated two-fold in M9 plus cholesterol (5 μg/ml). 5 ml aliquots were prepared and 1000 worms were added in a volume of 10-20 μl. NAC (500 mM stock solution in H_2_O) was added to a final concentration of 10 mM to half of the aliquots and incubated under rotation for 2 h. Thereafter, various concentrations of FA (10 mM and 12 mM) were added to the tubes and incubated under rotation for another 4 h. To this end, methanol-free 16 % FA (w/v; Thermo Scientific) was first adjusted to a 1 M stock solution in H_2_O, which was prepared fresh for each experiment. After the NAC/FA treatment, the solutions were centrifuged at 1300 rpm for 1 minute, followed by two washing steps with 5 ml M9 medium. Finally, worms were pelleted again and the volume was reduced to 500 μl. A volume of 25-50 μl (approx. 50-100 worms) was transferred to OP50-seeded NGM plates, on which the survival rate was scored under a stereoscope. Worms were qualified as dead when no locomotion could be detected and when stimulation with a wormpick did not cause a response. The survival count was repeated after 24 h, 48 h and 72 h. In parallel, developmental stages of worms were determined under the stereoscope at 48 h and 72 h post-treatment and qualified in the categories L1-L3, L4 and adult.

The combined PQ and FA treatment was performed in the same way as the NAC/FA treatment, with the exception that PQ was added at the same time as FA (2 mM) and incubated together for a total of 5 h. PQ (Methyl viologen dichloride hydrate, Sigma-Aldrich, #856177) was always freshly prepared and first adjusted to a 1 M stock in H_2_O, which was further diluted for the treatment.

#### Viability and survival assays

For determining cell viability, cells were seeded into 96-well plates at a density of 3000 cells per well and allowed to attach for 24 h at 37°C, 5 % CO_2_. Then, FA and/or L-buthionine-sulfoximine (L-BSO, Sigma-Aldrich, #B2515) and/or antioxidants were added to a final volume of 200 μl per well. 3 days later, resazurin (Sigma-Aldrich, #R7017) was added to a final concentration of 30 μM in the growing medium. Fluorescence (λ_ex_ = 525 nm; λ_em_ = 590 nm) was measured 3 h later in an Enspire Plate Reader (Perkin Elmer). For Nalm6, 5000 cells per well were seeded into 96-well plates and the drugs to be tested added immediately afterward. Viability was determined 5 days later. In all cases, the experiments were done by triplicate and data represented as percentage of the fluorescence obtained with the untreated samples of the corresponding cell line.

The colony survival assay was done by seeding 600 cells per well in 6-well plates. Immediately afterward, FA was added at the concentrations described in the text in a final volume of 2 ml (DMEM). Plates were maintained during 7-10 days at 37°C, 5 % CO_2_. Staining was done using a fixative/staining solution (0.5 % crystal violet, 6 % glutaraldehyde) for 30 minutes, following of extensive rinse with tap water. Visible colonies were counted, and the results expressed as percentage of the untreated wells. Experiments were done by duplicated and repeated the number of times indicated in each corresponding figure.

#### ROS measurement by H2DCFDA

ROS measurement was performed using 2’,7’-dichlorodihydrofluorescein diacetate (H2DCFDA, Sigma-Aldrich #D6883). Briefly, 4×10^4^ HCT116 cells were seeded per well in 24-well plates and allowed to adhere overnight. Cells were then treated with 0, 60 and 150 μM of FA and 0 or 100 μM of L-BSO, for 48 h. H_2_O_2_ 500 μM was used as a positive control and added 15 minutes prior H2DCFDA staining. After treatment, H2DCFDA was added to each well at a final concentration of 10 μM, kept 30 min at 37 °C. Then, cells were lifted and transferred to flow cytometry tubes, which were kept at 4°C until measuring was performed. Fluorescence (λ_ex_ = 488nm; λ_em_ = 530nm) was measured by flow cytometry using a Becton Dickinson’s FACS Canto II Flow cytometer.

#### ROS and GSH redox status determination by genetic sensors

The cytosolic roGFP2 sensor was cloned from Addgene 49435 into a retroviral backbone pLPCX by Gibson Assembly. Plasmid sequence was confirmed by sequencing. Retroviral infection was carried out transfecting HEK293T cells with pBS-CMV-gagpol (Addgene, #35614) and pCAG-VSVG (Addgene, #35616) vectors in addition to pLPCX cyto Grx1-roGFP2 (Addgene, #64975) or pLPCX cyto roGFP (Addgene, #49435). Conditioned medium was collected and recipient HCT116 cells infected adding Polybrene (Merk, #TR-1003-G) (1 μg/μL). Infection was confirmed by GFP-expression (90 % efficiency). Cells expressing the desired reporter were selected with 0.5 μg/mL puromycin. No clonal selection was carried out to prevent single clone artefacts. For ROS measurement, cells were seeded into a 24-well plate at 3.5×10^4^ cells per well and allowed to adhere overnight. Cells were then treated with 0, 60 and 150 μM of FA, and 0 or 100 μM of L-BSO, for 48 h. H202 500 μM, which was used as a positive control, was added 15 minutes prior cell analysis. After treatment, culture media was removed, and cells were washed with PBS. Cells were then trypsinized and transferred into clear flow cytometry tubes containing phosphate buffer saline (PBS) supplemented with 2 % fetal bovine serum (FBS). Tubes were kept at 4°C until measuring was performed. Fluorescence (λ_ex_ = 405nm and 488nm, λ_em_ = 510nm) was measured by flow cytometry using a Becton Dickinson’s FACS Aria II flow cytometer.

For determination of the glutathione (GSH) redox potential, cells expressing pLPCX cyto Grx1-roGFP2 were exposed to FA and/or L-BSO for 48 h. Then, cells were collected and fluorescence (λ_ex_ = 405nm and 488nm, λ_em_ = 510nm) determined by flow cytometry as described above^23^. The fraction of oxidized Grx1-roGFP2 sensor was calculated using the formula:

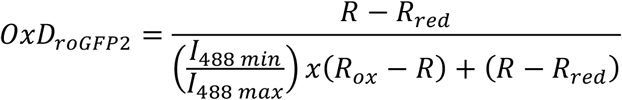

#### Metaphases analysis

To assess single-chromosome damage, HCT116 Wild type (WT) and Δ*ADH5* cells were plated in P60 dishes allowed to adhere and then treated with mitomycin C (MMC, Santa Cruz, #sc-3514) 0.5 μg/ml during 24 h or with FA 150 μM during 48 h. 16 h before harvesting the cells Colcemid (Gibco, #15212-012) was added at the concentration of 0.08 μg/ml without replacing the culture medium. Cells were washed with PBS and trypsin added to a final concentration of 0.125 %. Complete medium was added to stop trypsin reaction and clumps of cells disrupted by pipetting. Then, cells were centrifuged and resuspended into 2 ml of prewarmed hypotonic solution (KCl 0.075 mM) and incubated in 14 ml of this solution for 15 minutes at 37 °C. Then, 1 ml of fixative solution was added (3:1 methanol:glacial acetic acid) dropwise. Cells were washed twice with fixative solution and then dropped onto chilled humid slides, where cells were left to dry overnight. The day after, slides were stained in 2 % Giemsa solution (Thermo Scientific, #10092013) prepared in Gur buffer (Gibco, #10582-013), left to dry and mounted using BC solution (Cicarelli, #891). Pictures were taken using a Zeiss Axiobserver Z1 microscope with a 40x oil-immersion objective and analyzed using ImageJ^46^. To guarantee unbiased quantitation, pictures were taken by a microscopy technician, who labeled the images with numbers. After scoring of chromosome damage, the identities of the images were revealed.

#### 3D-Spheroid assay

96-well plates were coated with 50 μL of 1.5 % sterile agarose 2 h before cell plating. 100 μl of HCT116 cells (WT, Δ*ADH5*, Δ*ADH5* complemented, Δ*GCLM* or Δ*ADH5* Δ*GCLM*) were seeded at a concentration of 2×10^3^ or 4×10^3^ cells/well. Immediately after seeding, cells were treated with 100 μl of DMEM 10 % FBS containing 2x concentrations of the drugs used. The final concentrations of the drugs were 0, 50, 100 and 150 μM FA; 0 and 100 μM L-BSO; 0 and 500 μM NAC; 0, 1 mM Glutathione monoethyl ester (GSH-MEE, Santa Cruz, # sc-203974); 0 and 1 mM Trolox (Sigma-Aldrich, ##238813). Plates were kept at 37 °C. Spheroid formation was assayed by microscopy (Zeiss Axio A1 inverted microscope) 5-7 days after seeding and registered using a CANON Rebel T3i camera attached to the microscope with an appropriate adaptor, at 40x magnification. For sphere-size quantification Fiji was used to measure the area of the formed sphere.

#### Phylogenetic analysis

Eukaryote orthologs of ADH5 were obtained from NCBI, CLUSTAL at phylogeny.fr was used to align the sequences and TreeDyn at phylogeny.fr for tree generation^47^.

#### Western blot analysis

Cells were washed with PBS containing 1 mM N-Ethylmaleimide (NEM, Santa Cruz, #sc-202719), then lysed with Laemmli Sample buffer containing 2 % SDS, 4 % glycerol, 40mM Tris-Cl (pH 6.8), 5 % 2-Mercaptoethanol, 0.01 % bromophenol blue, 1 mM NEM, 1 mM Phenylmethylsulfonyl fluoride (PMSF), protease inhibitor mixture (Roche, #COEDTAF-RO), and phosphatase inhibitor mixture (Roche, #04906837001). Samples were bath-sonicated (3 pulses 30’’ ON 30’’ OFF) and boiled for 10 minutes. Sample concentration was relativized by Coomassie Brilliant Blue staining. For separation, samples were loaded onto 12 % polyacrylamide gels and subjected to electrophoresis. Protein was transferred to nitrocellulose membranes, which were blocked with 2 % BSA in Tris-buffered saline (TBS) or 5 % non-fat milk in TBS. Membranes were incubated with primary antibodies overnight at 4°C, followed by incubation with secondary antibodies conjugated with either horseradish peroxidase or fluorescent dye. DNA damaging agents were MMC, cisplatin (Santa Cruz, #sc-200896) and Hydroxyurea (Santa Cruz, #sc-29061) Proteins were visualized using ECL prime chemiluminescence reagent or fluorescence emission, respectively. Primary antibodies used were p53 (CST, #9282), phospho-P53 (CST, #9284), p21 (CST, #2947), phospho-histone H2A.X (CST, #9718), Vinculin (Santa Cruz, #sc-73614), alpha-tubulin (CST, #2144), and beta-actin (Santa Cruz, #sc-47778). Secondary antibodies used were horseradish peroxidase-linked anti-rabbit (CST, ##7074), horseradish peroxidase-linked anti-mouse (CST, #7076), DyLight-800 4x PEG-linked anti-rabbit (CST, #5151), and DyLight 680-linked anti-mouse (CST, #5470).

#### Cell cycle assay and apoptosis determination

Cells were plated at a final concentration of 3×10^5^ cells per well in DMEM supplemented with 10 % FBS. After 24 h, cells were treated with 0, 60, and 150 μM FA for 24 h. After this period, cells were harvested by trypsinization and pelleted by centrifugation (5 min, 1000 x g). Cells were washed with cold PBS and then fixed with 70 % cold ethanol for 15 min on ice. Cells were washed twice with PBS and treated with 30 μg ribonuclease A and 15 μg of propidium iodide. Cells were run on a BD FACS Canto II flow cytometer and the data was analyzed with FlowJo 10.0.7 (Tree Star). For apoptosis determination, the BD PE Annexin V Apoptosis Detection Kit (BD Pharmigen, #579563). Briefly, cells were plated and 24 h later exposed to the indicated concentrations of FA. 24 h later, cells were lifted, washed with cold PBS and stained with PE-Annexin V antibody and 7-AAD. Samples were run on a BD FACSAria II flow cytometer and data analyzed with FlowJo 10.0.7 (Tree Star).

#### GSH measurement

GSH was determined using the GSH-Glo™ Glutathione Assay (Promega, #V6911). Briefly, 10000 cells per well were seeded in a 96-well plate. A duplicated plate was prepared to determine viability. 48 h later GSH was determined following the instructions provided in the kit. In parallel, the viability was scored using resazurin and the results adjusted for the percentage of viable cells relative to the GSH content of WT cells.

#### Formaldehyde determination in blood

Mice were sacrificed at 6 months of age by decapitation, full blood was collected, and serum was separated from red blood cells by centrifugation (15,000 x g, 30 min, 4 °C). Serum was transferred to a new Eppendorf tube and sera of three mice of the same age, sex and diet were pooled and subsequently subjected to trichloroacetic acid (Guoyao, #80132618) (20 % w/v, in ultrapure water) precipitation. Therefore, trichloroacetic acid was added in a 1:1 ratio to the serum, vortexed for 30 sec. and centrifuged (15,000 x g, 30 min, 4°C). The supernatant was transferred to a fresh Eppendorf tube and stored at −80 °C until further processing.

The concentration of FA was detected by high-performance liquid chromatography (HPLC) as previously described^48^. Serum samples (0.08 ml each) were mixed with 0.02 ml 10 % trichloroacetic acid, 0.08 ml acetonitrile (Thermo Scientific, #A998-4), and 0.02 ml 2,4-dinitrophenylhydrazone (Beijingshiji, #550626). Samples were centrifuged (15,000 x g, 4 °C, 10 min) and then reacted in a 60°C water bath for 30 min; this step was followed by a centrifugation (15,000 x g, 4 °C, 10 min) and filtered (0.22 μm). 20 μl of the solution was then subjected to HPLC (LC-20A, Shimadzu, Japan). FA-DNPH derivatives were detected with an ultraviolet detector (Cas: 228-34016, Shimadzu, Japan) and a C18 reversed-phase column (Sigma-Aldrich, #50208-U), using 65 % acetonitrile as the mobile phase.

#### Synthesis of *S*-hydroxymethyl-glutathione

##### Reaction

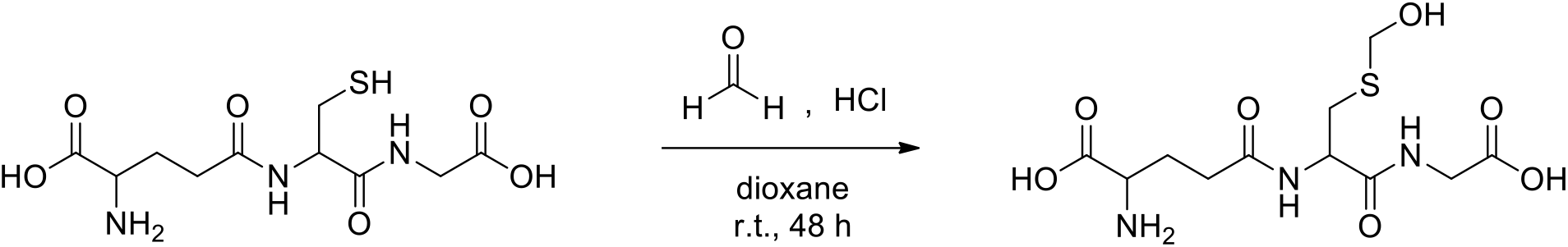

##### Procedure

In a 50 ml-bottom flask, FA solution (12 μL, 37 wt. % in water, Sigma-Aldrich, #F8775) and HCl (0.5 ml, 36.5-38.0%) were dissolved in 2 ml of dioxane (Sintorgan, #SIN-083003-63). The mixture was stirred at room temperature over 5 min and glutathione (Santa Cruz, #sc-29094) (50 mg, 0.16 mmol) was added in small portions. After stirring at room temperature over 48 h, the mixture was neutralized with saturated aqueous NaHCO_3_ solution and partitioned between ethyl acetate and water. The aqueous phase was lyophilized (0.03 mBar, −80 °C, 72 h) to obtain a white solid using a Telstar LYOQuest-85 freeze dryer (Telstar, Madrid, Spain). A portion of 10 mg of the solid was resuspended in 1 ml of a CH_3_OH:CH_3_CN (1:1) mixture, centrifuged and the supernatant was diluted to be analyzed by UPLC-HRMS. Estimated reaction yield 95.8 %^49^.

#### Sample preparation for UPLC-HRMS analysis

HCT116 WT and Δ*ADH5* cells were counted and cultured in 100 cm plates at 1×10^6^ cells/plate. Two independent rounds of sample preparation were carried out in consecutive weeks. 6 plates in the first week and 5 plates in the second week for each cell line were set up and allowed to grow for 72 h. One plate in each round was used for protein and cell count. Once 80 % confluence was reached, cells were gently washed with 5 ml of a 0.9% NaCl aqueous solution at 0 °C. Subsequently, enzymatic activity was quenched by adding liquid N_2_. Cells were scrapped immediately after with 1.4 ml of a cold (0 °C) CH_3_OH:CH_3_CN (50:50 v/v) solution and subsequently frozen using liquid N_2_. After one freeze-thaw cycle, samples were vortex-mixed during 30 s and centrifuged at 5000 × *g* for 5 min at 4 °C. Supernatants were collected and stored at −20 °C for 2 h and subsequently centrifuged at 15000 × *g* for 10 min at 4 °C. Afterwards, 1.4 ml of ultrapure water was added to supernatants and these solution were immediately frozen and stored at −80 °C until lyophilization.

Process blanks consisting of incubating culture media in plates without cells were generated in parallel with samples, and followed the same protocol described above. For protein and cell count, cells were lifted and counted using trypan blue as viability marker. Afterwards, cells were lysed in a solution containing 1 μM EDTA; 10 μM Tris pH 8; 200 μM NaCl and 0.2 % Triton, and total protein was determined by the Bradford assay using BSA as standard. Samples were lyophilized at −80 °C and 50 mTorr for 48 h using a Telstar LYOQuest-85 freeze dryer (Telstar, Madrid, Spain) and stored at −80 °C until analysis by UPLC-HMRS. All sample residues from each batch were reconstituted the same day in a water: methanol (90/10 v/v) solution. Reconstitution factors were selected to reach the same protein content for all samples. After reconstitution, samples were vortex-mixed for 30 s and centrifuged at 21382 x *g* for 20 min and 4 °C. Supernatants were stored until use at −80 °C. Quality control (QC) samples were prepared by pooling an aliquot of 15 μL from each sample, vortex-mixed for 30 s, split into 4 micro tubes, and stored at −80 °C until use for analysis.

A pooled QC sample spiked with GSH (14.3 μM), GSH disulfide (15.5 μM) and S-hydroxymethylglutathione (20 μM) was used to verify the stability of retention times, peak shapes and areas during the analysis.

#### UPLC-HRMS analysis

UPLC-HMRS analyses were performed using a Waters ACQUITY UPLC I Class system fitted with a Waters ACQUITY UPLC BEH C_18_ column (2.1×100 mm, 1.7 μm particle size, Waters Corporation, Milford, MA, USA, catalog #186002352), and coupled to a Xevo G2S QTOF mass spectrometer (Waters Corporation, Manchester, UK, SN: YDA 375) with an electrospray ionization (ESI) source operated in ESI positive ionization mode. The typical resolving power and mass accuracy of the Xevo G2S QTOF mass spectrometer were 32,000 FWHM and 0.3 ppm at *m/z* 556.2771, respectively. The mobile phase consisted of water with 0.1 % formic acid (Fisher Chemical, #F/1900/PB15) (mobile phase A) and methanol (Fisher Chemical A454-4, (UN 1230-CL3)) (mobile phase B). The flow rate was constant at 0.3 ml min^-1^, the elution gradient was set as follows: 0-1.6 min 0-0 % B; 1.6-2 min 0-20 % B; 2-6 min 20-70% B; 6-7 min 70-70 % B; 7-14 min 70-90 % B; 14-17.5 min 90-90 % B; 17.5-18 min 90-95 % B; 18-21 min 95-95 % B. After each sample injection, the gradient was returned to its initial conditions in 9 min (total run time was 30 min). The eluates from the analytical column were diverted by automatically switching the valve to waste, except for the elution window from 0 to 8 min. The column and autosampler tray temperatures were set at 35 and 5°C, respectively. The injection volume was 2 μL.

A solvent blank, which consisted of a water: methanol (90:10 v/v) solution, and a process blank were analyzed at the beginning and end of each batch. Samples were randomly analyzed within a defined template of spiked QC samples, and the analysis order was balanced based on sample classes. QC samples were used to condition the LC-MS system before sample analysis. A total of 20 randomized samples (WT cells n=10 and Δ*ADH5* cells n=10) were analyzed along 3 consecutive days. UPLC–MS sample lists were set up as follows (sample type (technical replicates)): zero consisting of mobile phase analysis without injection (1); solvent blank (2); process blank (2); QC samples (5); spiked QC sample (1); randomized, and balanced samples (12) with 1 spiked QC sample analyzed every 4 samples; spiked QC sample (1); process blank (2); solvent blank (1).

The mass spectrometer was operated in positive ion mode with a probe capillary voltage of 2.5 kV and a sampling cone voltage of 30.0 V. The source and desolvation gas temperatures were set to 120 and 300 °C, respectively. The nitrogen gas desolvation flow rate was 600 L h^-1^, and the cone desolvation flow rate was 10 L h^-1^. The mass spectrometer was daily calibrated across the range of *m/z* 50-1200 using a 0.5 mM sodium formate solution prepared in 2-propanol/water (90:10 v/v). Data were drift corrected during acquisition using a leucine encephalin (*m/z* 556.2771) reference spray (Waters cop, #700008842) infused at 5 μl min^-1^, every 45 seconds. Data were acquired in MS continuum mode in the range of *m/z* 50-1200, and the scan time was set to 0.5 seconds.

Principal component analysis (PCA) was conducted using MATLAB R2015a (The MathWorks, Natick, MA, USA) with the PLS Toolbox version 8.1 (Eigenvector Research, Inc., Manson, WA, USA). PCA was used to track data quality and to identify and remove outliers in the dataset. Two samples were identified as outliers by PCA, one from WT and one from Δ*ADH5* cells, and were not further considered for data analysis.

For UPLC-MS/MS experiments, the product ion mass spectra were acquired with collision cell voltages between 10 and 30 V, depending on the analyte. Ultra-high-purity argon (≥99.999%) was used as the collision gas. Data acquisition and processing were carried out using MassLynx version 4.1 (Waters Corp., Milford, MA, USA).

Chemical standards were prepared in ultrapure water and were analyzed under identical conditions as samples to validate metabolite identities by chromatographic retention time and MS/MS fragmentation pattern matching. Spiking experiments were also conducted with the authentic chemical standards on samples to address retention time differences caused by matrix effects.

Two different normalization strategies were independently used for sample analysis: data were normalized by number of viable cells or by protein content.

#### Quantification and Statistical Analysis

Prism software package (GraphPad Software 7) was used for statistical analysis with the level of significance of 0.05 (95 % confidence) and one-way ANOVA using the Tukey’s algorithm for multiple comparisons. For mass spectrometry data whisker plots and Mann-Whitney test was used to asses significance. The box and whiskers plots are represented by a line in the box corresponding to the median; the edges are the 25th and 75th percentiles and the whiskers extend to the most extreme values in data. Additional information about statistical tests, sample number and P-values are described in figure legends. Unless otherwise stated, experiments were done using technical replicates (2 or 3 wells per condition) and repeated the n times described in the figure legends with each symbol in a bar plot representing the average of the technical replicates for a given biological sample.

## Data availability

All data generated during this study are included in the published paper including source data for figure 2, Extended data Fig. 1 and Extended data Fig. 5.

## Acknowledgements

This work was supported by CONICET (PUE 2016 22920160100010CO), FOCEM MERCOSUR (COF 03/11), ANPyCT (PICT-PRH 2017-4668) and (PRH-PICT-2015-0022). LBP is a collaborative Group Leader between IBioBA, MPI for Metabolism Research (Cologne, Germany) and MPI for Biophysical Chemistry (Göttingen, Germany), from whom he receives support. B.S. acknowledges funding from the Deutsche Forschungsgemeinschaft (SCHU 2494/3-1, SCHU 2494/7-1, SCHU 2494/10-1, SCHU 2494/11-1, SFB 829, KFO 286, KFO 329, GRK 2407) and the Deutsche Krebshilfe (70112899). IK acknowledges EMBO (ALTF 1029-2014). LBP would like to acknowledge Prof. Jens Brüning and Prof. Thomas Wünderlich for providing support for mouse breeding at MPI-MR. Also, to IBioBA staff for general support, to Gerry Crossan, Ricardo Biondi, Alejandro Leroux and Carolina Perez Castro for the critical reading of the manuscript. CU, AM, HR, MM and AF are CONICET fellows. MEM, MRM, MB and LBP are Research Staff Members from CONICET.

## Author contributions

CU, AM, MS and HR carried out cellular experiments assisted by LBP. MR, KK and BS contributed with *C. elegans* data generation and analysis. MRM and MEM designed the sample preparation protocol for metabolite extraction, developed the UPLC-QTOF-MS-based method, and performed data analysis. MRM conducted UPLC-MS/MS experiments. AF and MB synthesized S-hydroxymethyl-GSH. AV and IK processed mouse blood samples. YW and RH measured blood FA. LBP conceived the work and wrote the paper. All authors revised the manuscript.

## Ethics declarations

The authors declare no competing interests.

## Supplemental Information titles and legends

**Extended Data Fig. 1.**
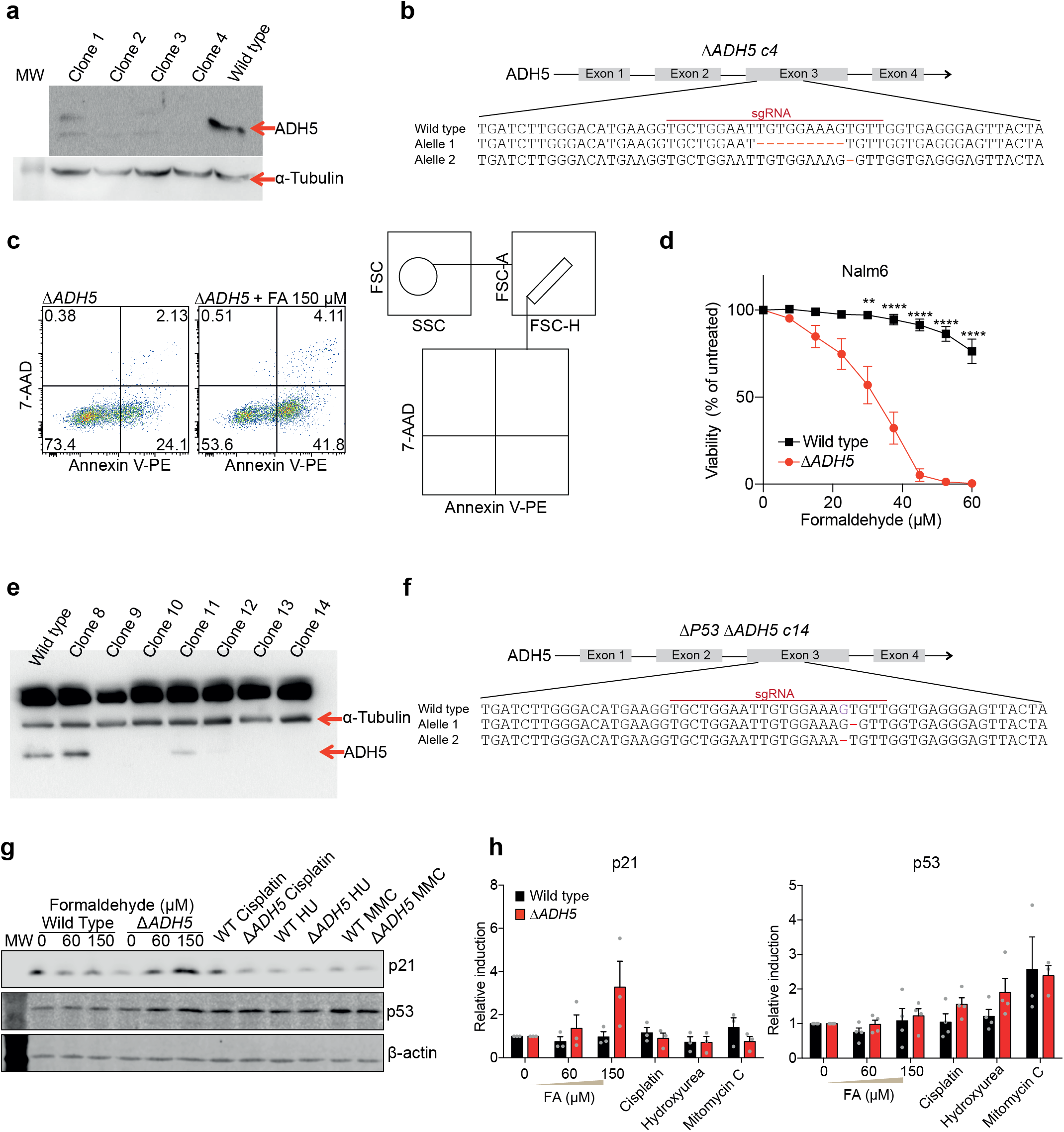
Δ*ADH5* cell generation and P53 response to formaldehyde. a. Western blot analysis of ADH5 expression in clones edited by CRISPR/Cas9. **b.** *ADH5* gene showing the exon targeted by CRISPR/Cas9 and the genetic modifications of the Δ*ADH5* clone used in this work. **c.** Left: Representative flow cytometry plot for Annexin V determination in Δ*ADH5* cells untreated or exposed to 150 μM formaldehyde (FA). Right: Gating strategy for determining apoptotic cells. **d.** Resazurin-based viability assay for Nalm6 cells exposed to increasing concentrations of FA (mean ± s.e.m., n=6, asterisks represent the statistical significance according to one-way ANOVA for multiple comparison using a Tukey-corrected testbetween WT and Δ*ADH5)*. **e.** Western blot analysis of ADH5 expression in Δ*P53* clones edited by CRISPR/Cas9. **f.** *ADH5* gene showing the exon targeted and the genetic modifications of the Δ*P53 ΔADH5* clone used in this work. **g.** Western blot against p21 and p53. Loading control β-actin. **h**. Quantitation of western blots against p21 (left) and p53 (right) (s.e.m., n=3).

**Extended Data Fig. 2.**
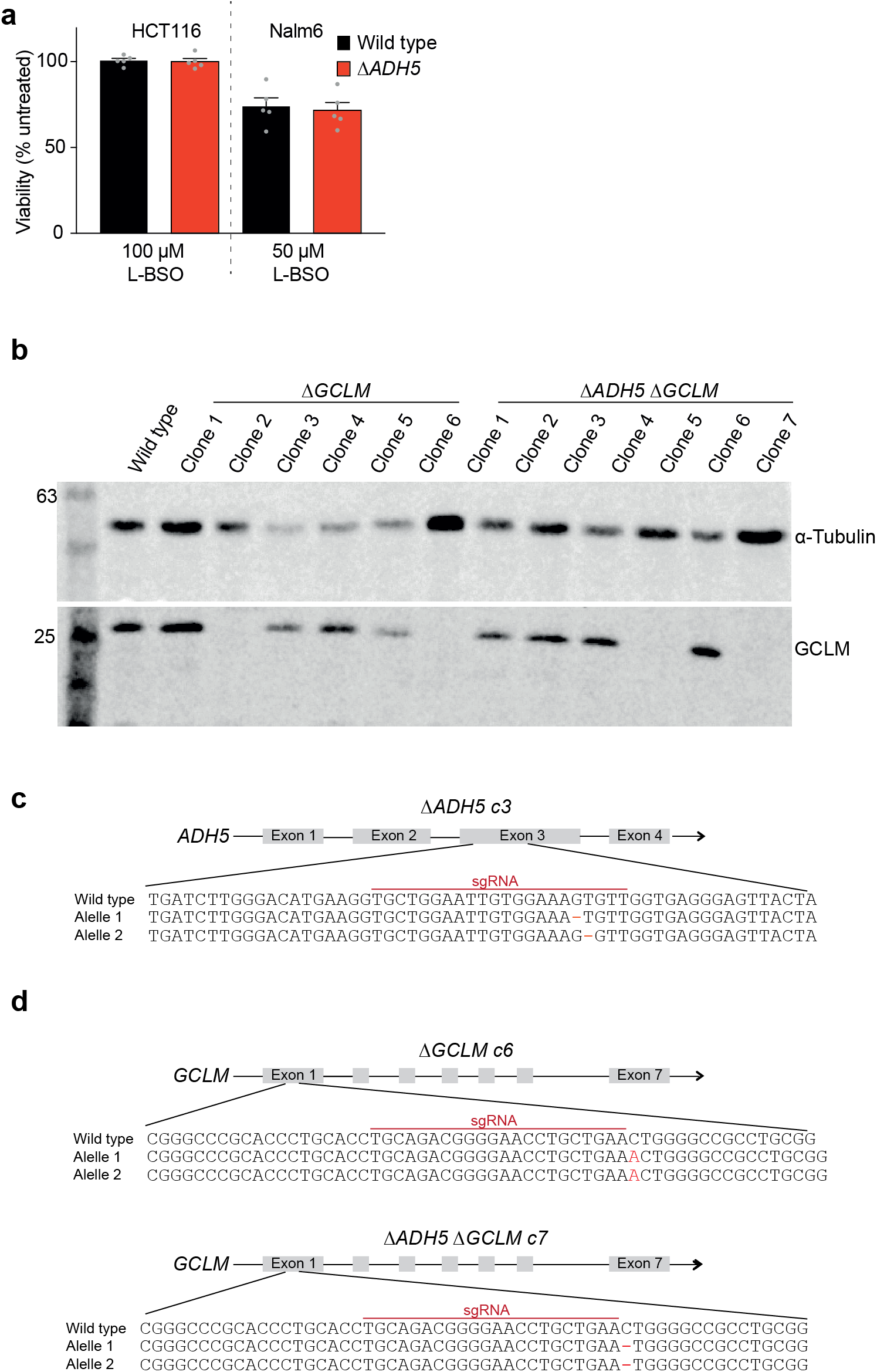
GSH biosynthesis inactivation. **a.** Resazurin-based viability assay for HCT116 (left) and Nalm6 (right) Wild type (WT) and Δ*ADH5* cells at 100 and 50 μM L-buthionine-sulfoximine (L-BSO), respectively (s.e.m., n=5). **b.** Western blot analysis of *GCLM* expression in clones edited by CRISPR/Cas9. **c.** *ADH5* gene showing the exon targeted by CRISPR/Cas9 and the genetic modifications of the Δ*ADH5* clone on which *GCLM* was inactivated. **d.** *GCLM* gene showing the exon targeted by CRISPR/Cas9 and the genetic modifications of the Δ*GCLM* clones used in this work.

**Extended Data Fig. 3.**
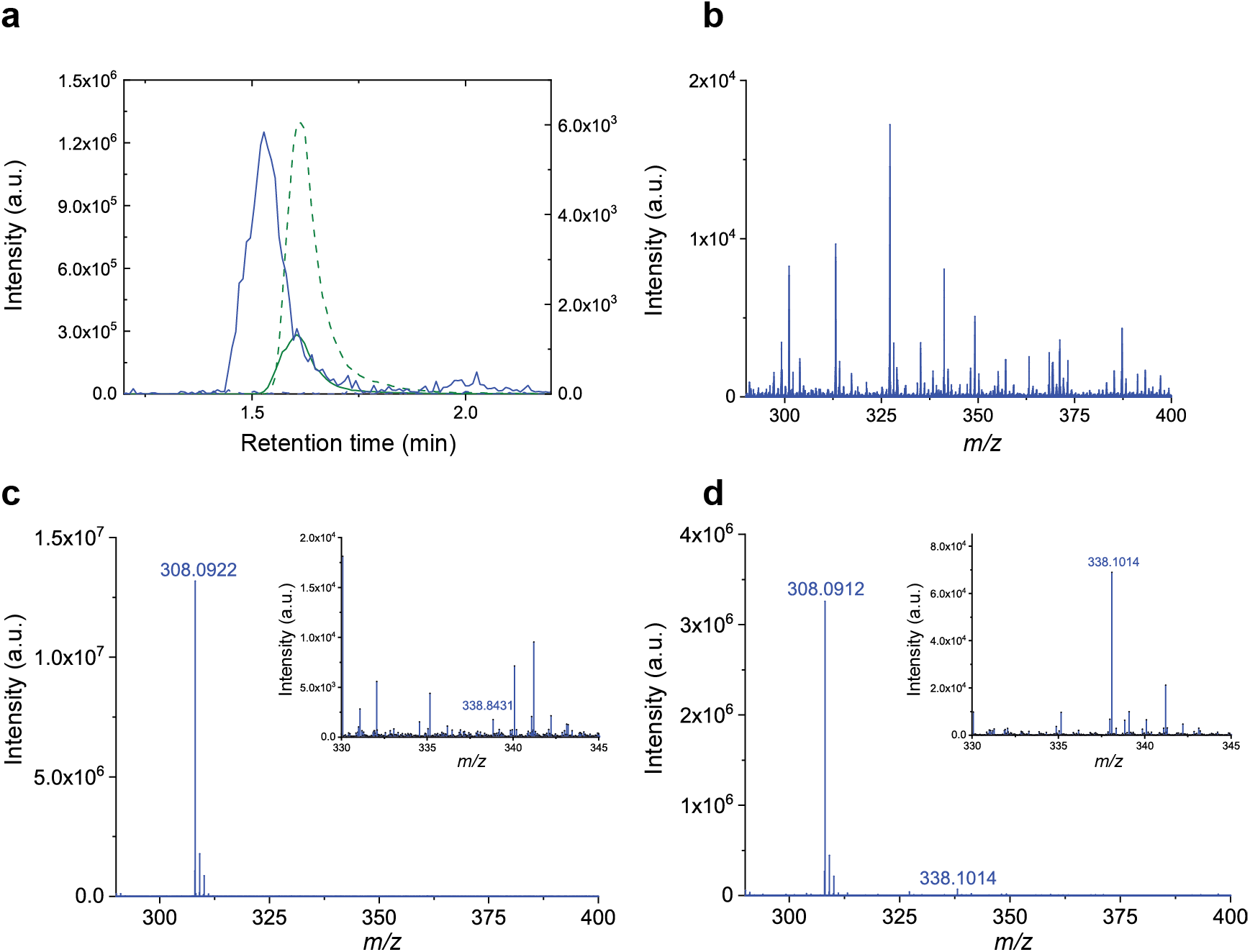
S-hydroxymethylglutathione synthesis. **a.** Extracted ion chromatograms for [GSH + H]^+^ ion at *m/z* 308.0916 generated from a 10.4 μM glutathione (GSH) standard solution before reaction (t_0_: green dash line) and after 48h of reaction (t_48_: green solid line); and for [S-hydroxymethylglutathione (HSMGSH) + H]^+^ ion at m/z 338.1022 generated from a 10.4 μM GSH standard solution before reaction (t_0_: blue dash line) and after 48h reaction (t_48_: blue solid line). **b.** Mass spectrum for the solvent at t_0_, with no signals detected at *m/z* 308.0916 or *m/z* 338.1022. **c.** Mass spectrum for a GSH standard solution at t_0_, with no signal detected at *m/z* 338.1022. **d.** Mass spectrum for the reaction mixture at t_48_.

**Extended Data Fig. 4.**
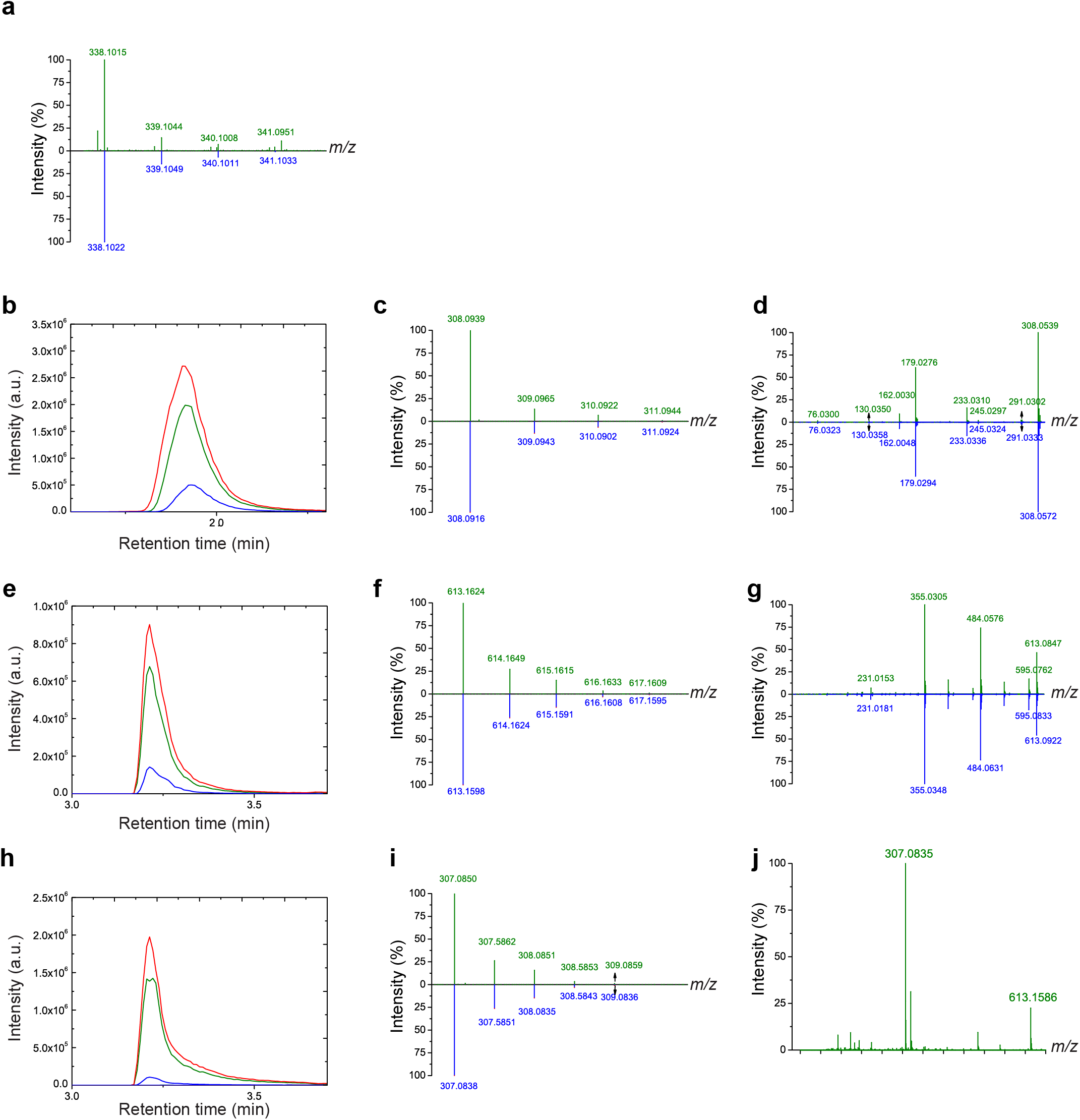
*in vivo* detection of S-hydroxymethylglutathione. **a.** Mass spectrum for [S-hydroxymethylglutathione (HSMGSH) + H]^+^ ion at *m/z* 338.1022 in a Wild type (WT) sample (green), and its simulated isotopic pattern (blue). **b.** Extracted ion chromatograms for [Glutathione (GSH) + H]^+^ ion at *m/z* 308.0916 ± 0.0500 generated from a non-spiked QC sample (green), a 43 μM spiked QC sample (red), and a 14.3 μM GSH standard solution (blue). **c.** Mass spectrum for [GSH + H]^+^ ion at *m/z* 308.0916obtained from a QC sample (green), and its simulated isotopic pattern (blue). **d.** Product ion mass spectrum for [GSH + H]^+^ precursor ion obtained from a QC sample (green), and a 14.3 μM GSH standard solution (blue), using a collision cell voltage of 10 V. **e.** Extracted ion chromatograms for [GSH disulfide(GSSG) + H]^+^ ion at *m/z* 613.1598 ± 0.0500 generated from a non-spiked QC sample (green), a 15.5 μM spiked QC sample (red), and a 15.5 μM GSSG standard solution (blue). **f.** Mass spectrum for [GSSG + H]^+^ ion at *m/z* 613.1598 obtained from a QC sample (green), and its simulated isotopic pattern (blue). **g.** Product ion mass spectrum for [GSSG + H]^+^ precursor ion obtained from a QC sample (green), and a 15.5 μM GSSG standard solution (blue) using a collision cell voltage of 20 V. **h.** Extracted ion chromatograms for [GSSG + 2H]^2+^ ion at *m/z* 307.0838 ± 0.0500 generated from a non-spiked QC sample (green), a 15.5 μM spiked QC sample (red), and a 15.5 μM GSSG standard solution (blue). **i.** Mass spectrum for [GSSG + 2H]^2+^ ion at *m/z* 307.0838 obtained from a QC sample (green), and its simulated isotopic pattern (blue). **j.** Mass spectrum for [GSSG + H]^+^ precursor ion obtained from a 15.5 μM GSSG standard solution (blue).

**Extended Data Fig. 5.**
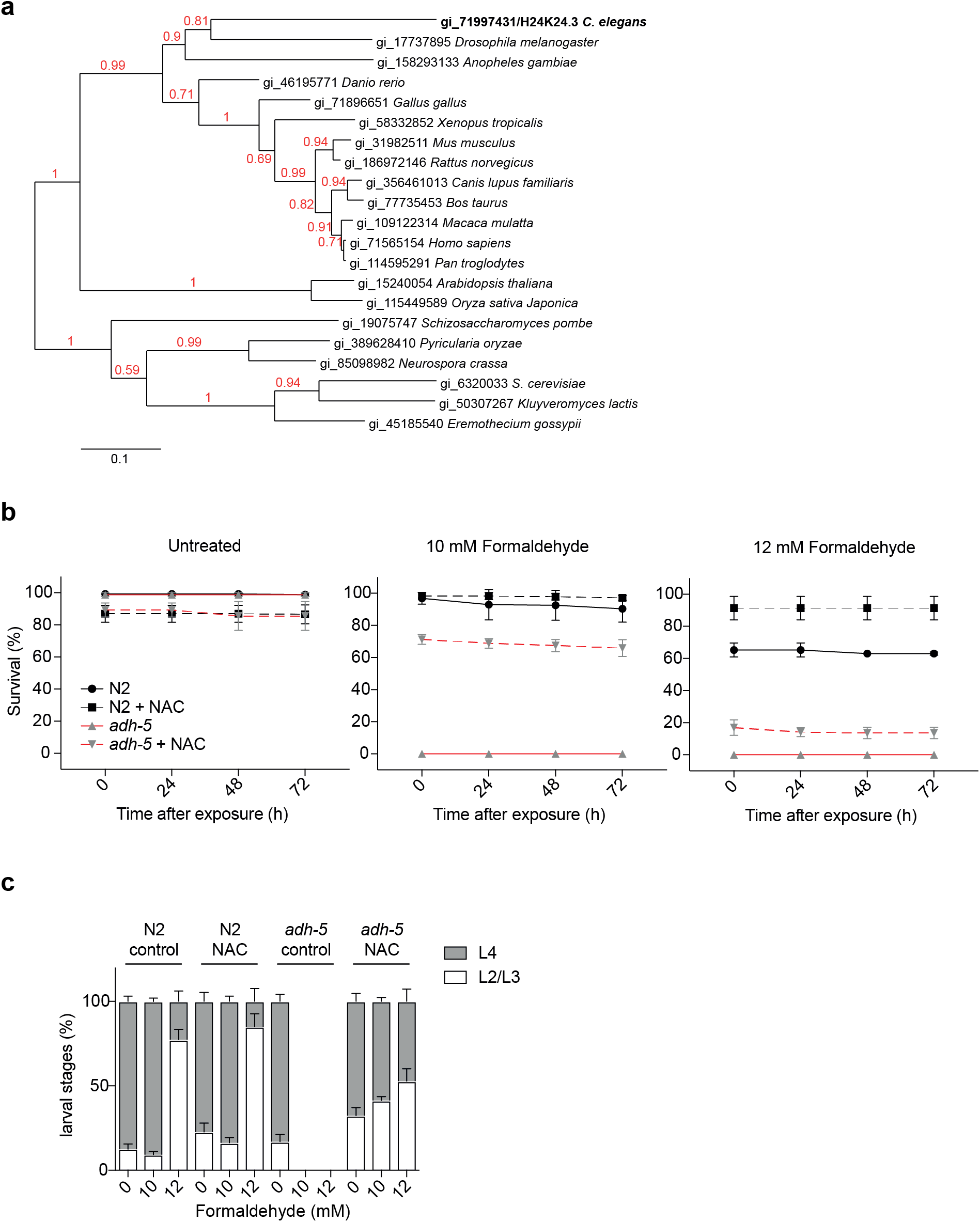
ADH5 is conserved and prevents FA toxicity in *C. elegans*, related to Figure 6. **a.** Phylogenetic analysis of *ADH5*-homolog genes in eukaryote highlighting the ortholog gene (gi_71997431/H24K24.3) found in *C. elegans*. **b.** Survival of L1-staged Wild type (N2) and *adh-5* mutant upon exposure to the indicated FA concentrations and 10 μM NAC measured 0, 24, 48 and 72 h after treatment (mean ± S.D., n=3). **c.** Development profile of surviving animals 48 h after FA exposure (mean ± S.D., n=3).

## Source data

**Source Data Fig. 2:**
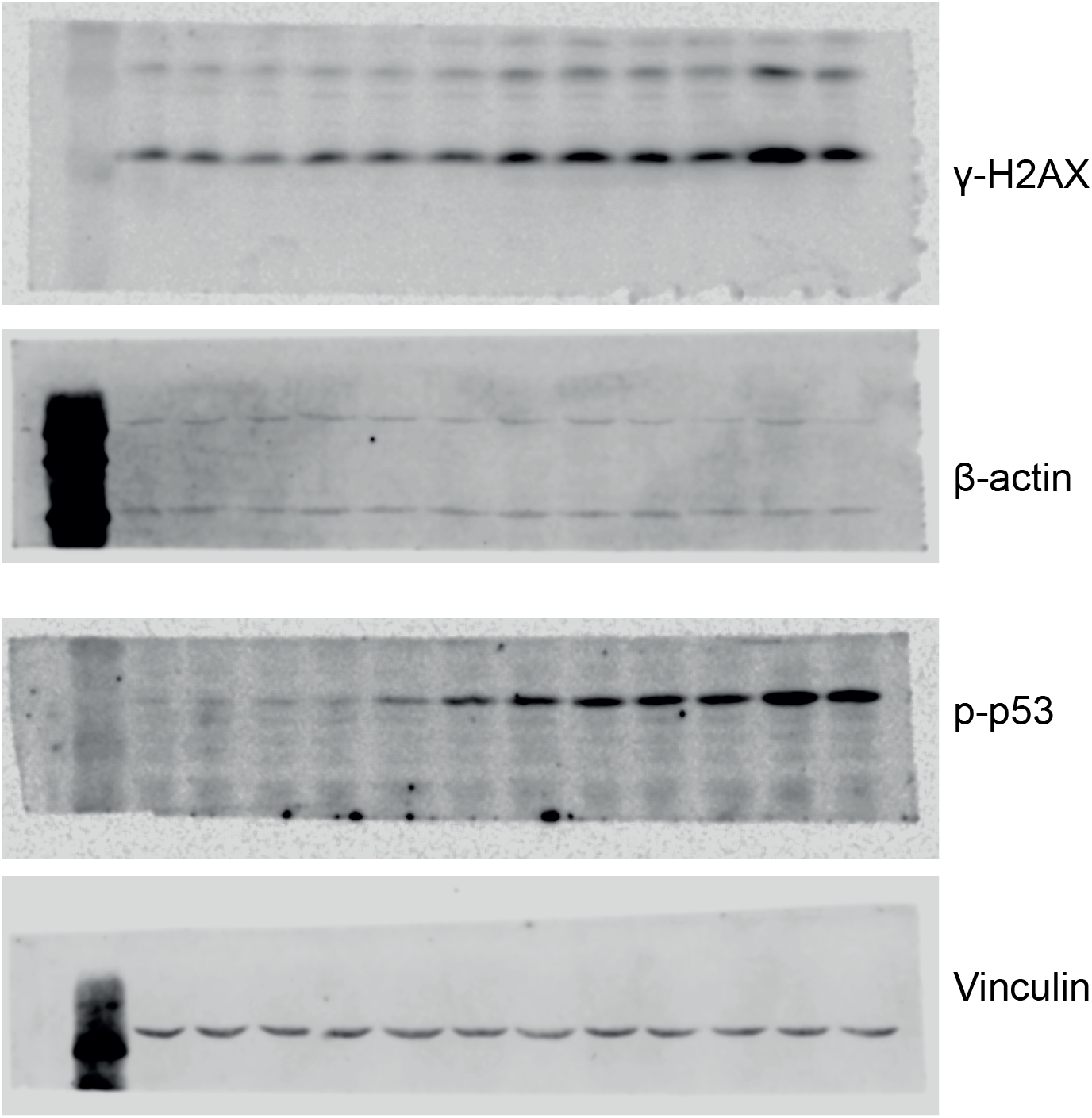
Uncropped images for western blots from Fig. 2

**Source Data Extended Data Fig. 1:**
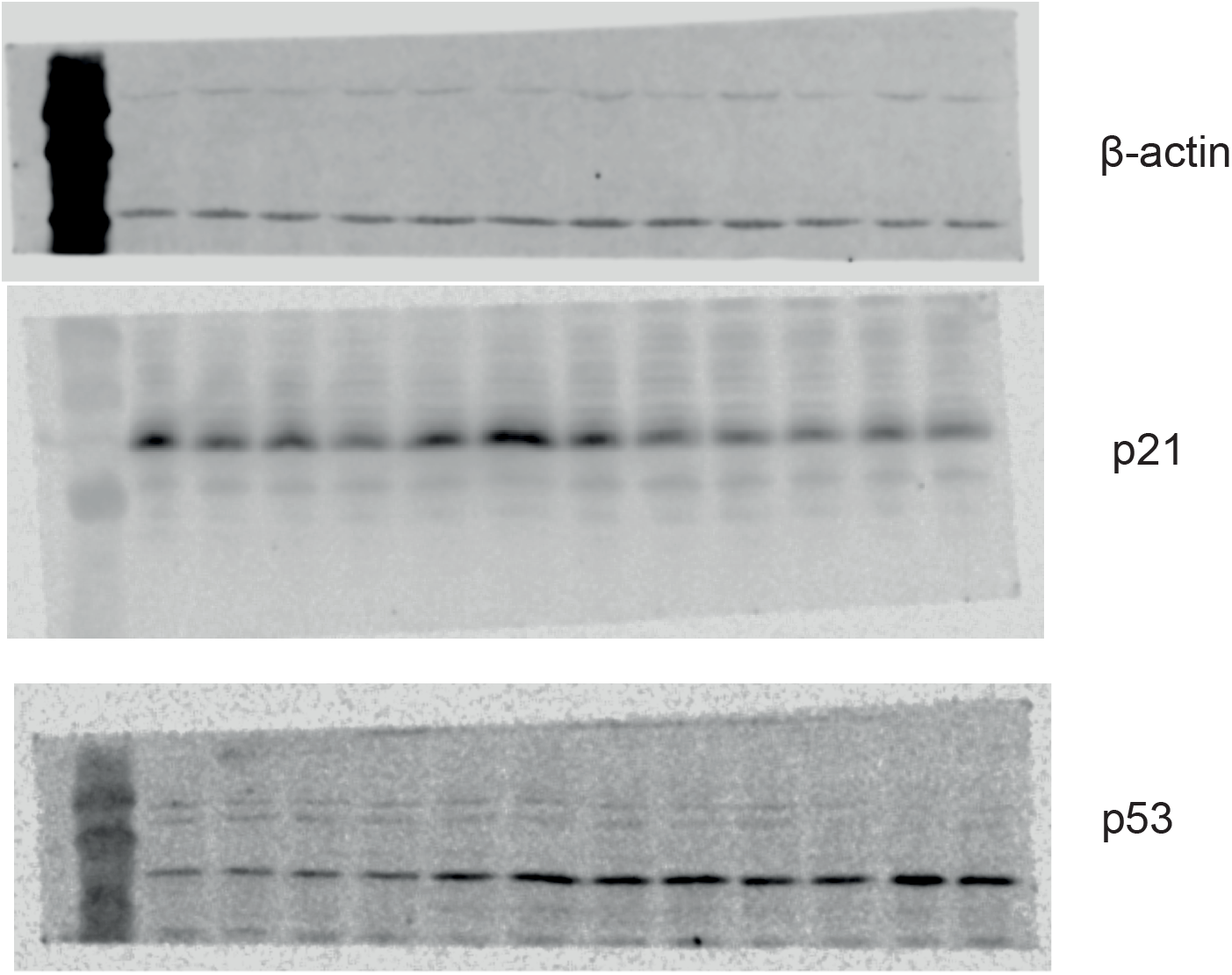
Uncropped images for western blots from Extended Data Fig. 1

**Source Data Extended Data Fig. 5.**
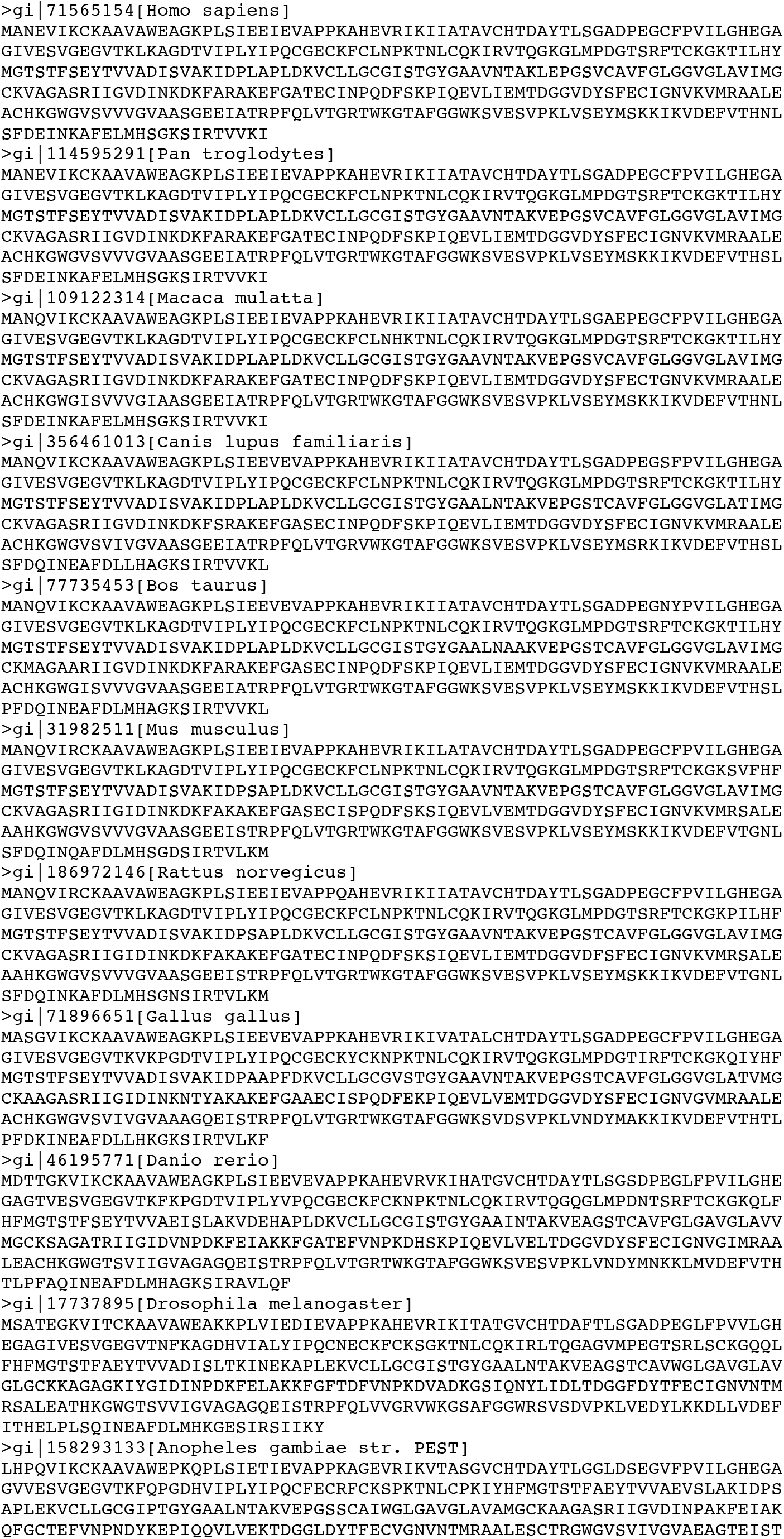
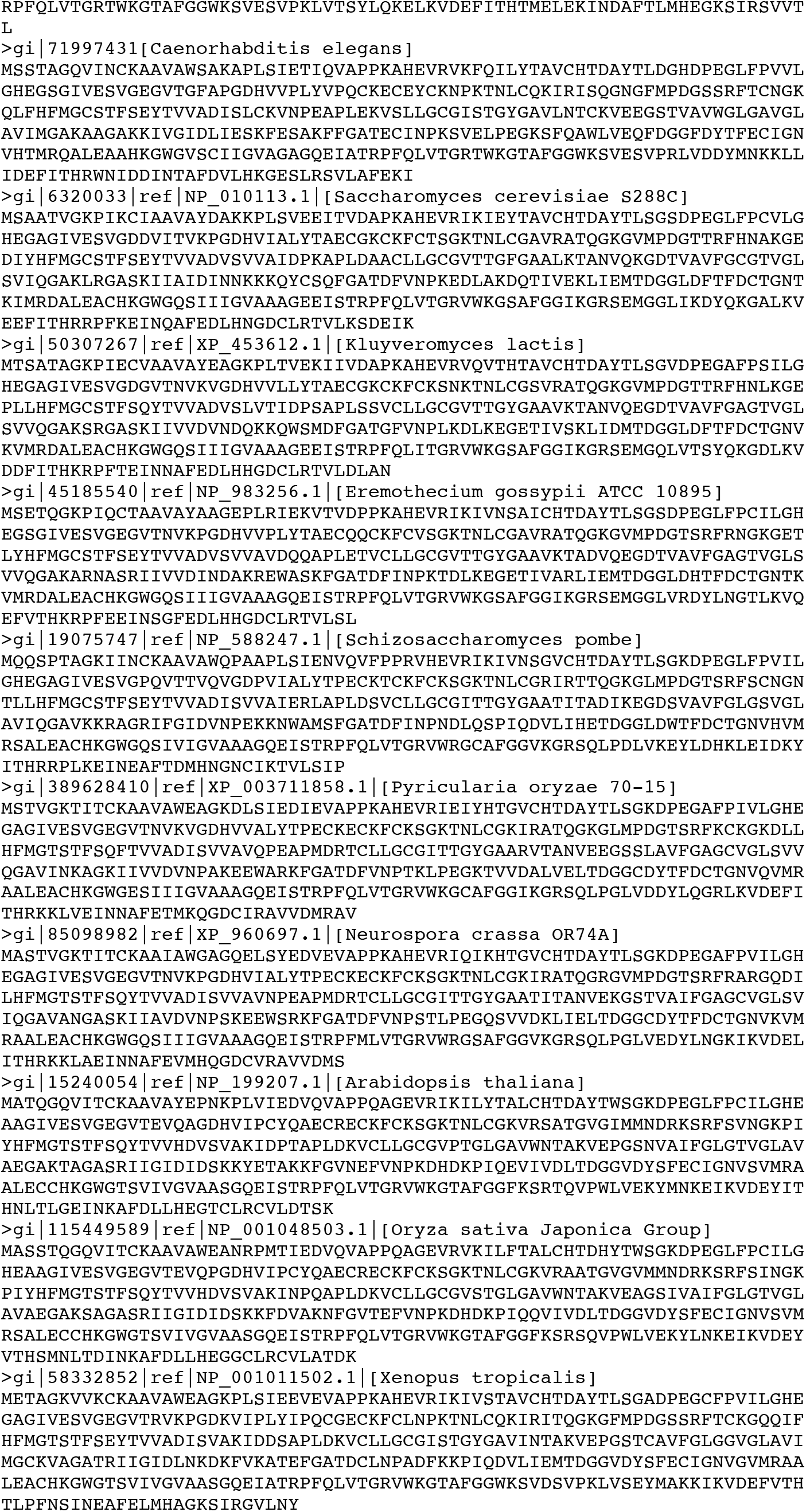
List of sequences used for phylogenetic analysis.

